# Machine learning to identify clinically relevant *Candida* yeast species

**DOI:** 10.1101/2023.09.08.556885

**Authors:** Shamanth A. Shankarnarayan, Daniel A. Charlebois

## Abstract

**Background:** Fungal infections, especially due to *Candida* species, are on the rise. Multi-drug resistant organism such as *Candida auris* are difficult and time consuming to identify accurately.

Machine learning is increasingly being used in health care, especially in medical imaging. In this study, we evaluated the effectiveness of six convolutional neural networks (CNNs) to identify four clinically important *Candida* species.

**Materials and Methods:** Wet-mounted images were captured using bright field live-cell microscopy followed by separating single cells, budding cells, and cell group images which were then subjected to different machine learning algorithms (custom CNN, VGG16, ResNet50, InceptionV3, EfficientNetB0, and EfficientNetB7) to learn and predict *Candida* species.

**Results:** Among the six algorithms tested, the InceptionV3 model performed best in predicting *Candida* species from microscopy images. All models performed poorly on raw images obtained directly from the microscope. The performance of all models increased when trained on single and budding cell images. The InceptionV3 model identified budding cells of *C. albicans, C. auris*, *C. glabrata* (*Nakaseomyces glabrata*), and *C. haemulonii* in 97.0%, 74.0%, 68.0%, and 66.0% cases, respectively. For single cells of *C. albicans, C. auris*, *C. glabrata*, and *C. haemulonii* InceptionV3 identified 97.0%, 73.0%, 69.0%, and 73.0% cases, respectively. The sensitivity and specificity of InceptionV3 were respectively 77.1% and 92.4%.

**Conclusion:** This study provides proof of concept that microscopy images from wet mounted slides can be used to identify *Candida* yeast species using machine learning quickly and accurately.

## Introduction

Fungi cause serious infections affecting more than one billion people of all ages across the globe and are responsible for approximately 1.6 million deaths per year ^1^. *Candida* species is the fourth leading agent to cause hospital acquired blood stream infections in USA ^2^, seventh in Europe ^3^, and affects 250,000 people worldwide and causes 50,000 deaths every year ^4^. *Candida* species, commonly found as commensals in human skin and mucosal surfaces, cause opportunistic infections when the immune system is impaired ^5^. The severity of the infection depends on the immune status of the individual, the infecting agent, and their virulence factors along with their ability to evade host factors ^6^. The mortality rate due to candidemia ranges from 22-75% ^7^.

Nearly 90% of invasive *Candida* infections are caused by *Candida albicans, Candida glabrata* (which has an alternate taxonomic name of *Nakaseomyces glabrata* ^8^)*, Candida tropicalis, Candida parapsilosis,* and *Candida krusei* ^9^. Due to the limited antifungal treatment options to treat invasive infections ^10^, early diagnosis and prompt antifungal therapy are critical for successful clinical outcomes. *Candida auris* has emerged as a multi-drug resistant pathogen causing hospital acquired infections and outbreaks. First reported in Japan in 2009, *C. auris* spread across the globe and recently the World Health Organization declared this pathogen as a “critical priority” pathogen ^11–13^. *C. auris* is commonly encountered in prolonged hospitalized patients and known to cause superficial to invasive blood stream infections ^14^. The mortality rate due to this pathogenic yeast ranges from 30% to 60% ^15^. In some hospitals in India, *C. auris* is the most isolated *Candida* species from blood culture ^16^. Previous studies indicate that *C. auris* exhibit increased minimum inhibitory concentrations (MICs) to all three major classes of antifungal drugs to treat invasive infections (i.e., azoles, polyenes, and echinocandins) ^17^. Several studies have reported that *C. auris* is often misidentified as *C. haemulonii*, *C. famata*, *C. sake*, and *Saccharomyces cerevisiae* by conventional laboratory methods (e.g. germ tube and carbohydrate fermentation tests) as well as automated commercial methods (e.g., VITEK and matrix assisted laser desorption ionization-time of flight mass spectroscopy (MALDI-TOF MS)) ^18–20^. Diagnostic-based antifungal treatments can reduce costs and shorten hospital stays compared to empirical treatments ^21^. Therefore, early detection of pathogenic *Candida* species can help improve patient outcomes and contain infectious fungal diseases.

Another major problem in the diagnosis of candidiasis is the delay in turnaround time from conventional methods. A few studies have utilized molecular techniques to reduce the turnaround time for the diagnosis of *Candida* infections. For example, Maaroufi et al. ^22^ developed a real-time PCR-based assay for the early detection and identification of commonly encountered *Candida* species from simulated blood cultures. This method showed high efficiency as an adjunct to blood culture systems with a short turnaround time (1-2 hours) compared to traditional methods. Farina et al. ^23^ evaluated the performance of MALDI-TOF MS in identifying clinically relevant *Candida* species directly from positive blood cultures. They found that MALDI-TOF MS had a prediction accuracy of 90% compared to conventional methods, with a much shorter turnaround time (12-24 hours). In addition to molecular methods, other techniques such as peptide nucleic acid fluorescence in situ hybridization (PNA-FISH) ^24^, flow cytometry ^25^, and Raman spectroscopy ^26^ have been explored in the context of rapidly identifying *Candida* species. However, these methods require sophisticated instruments (e.g., MALDI TOF mass spectrometer, DNA sequencer, etc.), extensive laboratory expertise, or time-consuming preprocessing of *Candida* species.

The “gold standard” for the diagnosis of the invasive candidiasis has been elaborated by Mycoses Study Group Education and Research Consortium group ^27^. In their consensus guidelines, four tests are stated to confirm the diagnosis of invasive candidiasis: histopathology findings, positive culture from sterile site samples, detection of yeast from paraffin-embedded tissue samples, and blood culture positivity (most employed test). Other non-culture methods employed for the diagnosis of the invasive candidiasis are beta-D-glucan ^28^, *Candida albicans* germ tube antibody ^29^, nucleic acid amplification test from blood samples^30^, and T2 Biosystems (*Candida* panel)-based diagnostic tests ^31^. However, these non-culture methods require expensive equipment and extensive expertise, limiting their usage in underdeveloped countries and in rural settings ^32^. Identifying these infecting *Candida* species is crucial due to high diversity among the genus and the possibility of intrinsic and acquired resistance in some of these species ^33^. The major drawback of these tests is their long turnaround times, which is approximately 72-96 hours, leading to delays in antifungal treatment resulting in increased mortality ^34^. Antifungal susceptibility testing takes an additional 48-72 hours to provide the susceptibility profile of the infectious agent, further delaying antifungal therapy.

Artificial Intelligence (AI) and machine learning (ML) are increasingly being applied in health care, especially in medical imaging for diagnosing diseases ^35, 36^. Images acquired via ultrasound, computed tomography, and magnetic resonance imaging, are being subjected to AI-computer aided design (CAD) for diagnostic purposes in clinical practice. AI-CAD has shown superior or similar performance compared to conventional diagnostic approaches for various diseases including ophthalmic diseases, respiratory diseases, and cancers ^37^. Deep learning is a form of AI with the capability to increase the accuracy as well as rapidity of the diagnosis through processing large medical image datasets, which has been deemed impossible for human experts^38^. Convolutional neural networks (CNNs) can classify images and detect objects in images ^39^. Deep learning has been used in infectious disease surveillance, including in early-warning systems for disease surveillance ^41^, pathogen classification ^42^, Kirby-Bauer disk diffusion assay interpretation (for bacteria) ^43^, nosocomial outbreak source identification ^44^, and risk assessment ^45^.

A limited number of studies have evaluated ML algorithms to identify yeast species. Data generated from the Raman spectroscopy were used to train ML models to identify *Candida* species ^26, 46^. Another study used stained *C. albicans* images from cosmetic products to check for contamination ^47^. Deep learning techniques have also been employed to capture *C. albicans* morphologies and explore their intermixing patterns ^48^. Other studies have combined technologies such as PCR, mass spectroscopy, molecular beacons, and laser induced break-down spectroscopy with machine to rapidly identify yeast species ^49–52^. These tests may offer improved “sensitivity” and “specificity” ^53^ (see “Definitions” section in supplementary materials) compared to traditional methods but require further validation before widespread implementation.

In this study, we evaluate the efficiency of ML models to identify four yeast species, namely *C. albicans*, *C. auris, C. glabrata* (*N. glabrata*), and *C. haemulonii*. Wet mounted images are captured by bright-field live-cell microscopy. These microscopy images are then separated into single cell, budding cell, and cell group images, which are used to train and validate the ML models to identify *Candida* species of pathogenic yeast. Specifically, we train, validate, and test deep neural network models to learn hierarchical representations from imaging data by stacking multiple layers of non-linear transformations. Overall, we find that ML models (especially the InceptionV3 model) can extract relevant features from the microscopy images to identify these yeast species.

## Material and Methods

### Strains, Media, and Growth Conditions

The authors confirm that the ethical policies of the journal, as noted on the journal’s author guidelines page, have been adhered to and the appropriate ethical review committee approval has been received. The US National Research Council’s guidelines for the Care and Use of Laboratory Animals were followed. *C. albicans*, *C. auris, C. glabrata*, and *C. haemulonii* isolates were obtained from clinical samples from the Alberta Precision Laboratories (APL)— Public Health Laboratory (ProvLab). All strains and isolates (**Table 1**) were preserved in 25% glycerol at -80°C until further use. The strains and isolates were revived by culturing from frozen stock on Sabouraud dextrose agar (SDA) plates (Millipore, Dramstadt, Germany) and incubated at 35°C for 48h. Fresh subcultures were made on SDA agar plates and incubated at 35°C for 24h prior capturing high-resolution microscopy images (see “Data collection” section in supplemental materials).

**Table 1:** Clinical *Candida* species isolates used in this study and their GenBank accession numbers.

| Species Number | Isolate | GenBank Accession Number |
| --- | --- | --- |
| 1 | *C. albicans* #1 | OR490529 |
| 2 | *C. albicans* #2 | OR490530 |
| 3 | *C. albicans* #3 | OR490531 |
| 4 | *C. albicans* #4 | OR490532 |
| 5 | *C. albicans* #5 | OR490533 |
| 6 | *C. auris* #1 | OP984814 |
| 7 | *C. auris* #2 | OP984815 |
| 8 | *C. auris* #3 | OP984816 |
| 9 | *C. auris* #4 | OP984817 |
| 10 | *C. auris* #5 | OP984818 |
| 11 | *C. glabrata* #1 | OR490534 |
| 12 | *C. glabrata* #2 | OR490535 |
| 13 | *C. glabrata* #3 | OR490536 |
| 14 | *C. glabrata* #4 | OR490537 |
| 15 | *C. glabrata* #5 | OR490538 |
| 16 | *C. haemulonii* | OR491713 |

### DNA Extractions, PCR, and Sequencing

All the clinical isolates were identified using MALDI TOF MS ^54^ by the APL-ProvLab. Then internal transcribed spacer (ITS) region of ribosomal DNA sequencing was carried out to confirm the *Candida* species identity. The primers ITS-5 (5’-GGAAGTAAAAGTCGTAACAAGG-3’) and ITS-4 (5’-TCCTCCGCTTATTGATATGC 3’) were used to amplify the ITS region (Integrated DNA Technologies, Iowa, USA). Genomic DNA extraction was done using the phenol-chloroform-isoamyl alcohol method as previously described ^55^. The quality and quantity of the extracted DNA was measured using a microvolume µDrop Plate (Thermo Fisher Scientific, Mississauga, Canada). The template and the primers were mixed in concentrations of 7.5ng/µL and 0.25µM, respectively, to a final volume of 10µL. Sanger sequencing was then performed using a 3730 Genetic Analyzer (Thermo Fisher Scientific, Mississauga, Canada, RRID:SCR_018052) at the Molecular Biology Services Unit at the University of Alberta. The resulting sequences were subjected to nucleotide BLAST analysis^56^ (RRID:SCR_001598), which revealed 100% similarity to the standard strains. The isolates’ ITS sequences were submitted to NCBI with the accession number OR490529-OR490538 (**Table 1**).

### Data Collection

The isolated colonies of different *Candida* species grown on SDA medium for 24h was utilized to prepare wet mount using normal saline (0.9%) on the microscopic slide (Fisherbrand, Pittsburg, USA) overlayed with the coverslips (size: 22mm, Fisherbrand, Pittsburg, USA). Raw images (1000 images) for each species were captured using EVOS M7000 imaging system (Invitrogen, Thermo Fisher Scientific, Massachusetts, USA) at 100x resolution. Coarse and fine adjustments were made to focus the cell on a single plane before capturing the images. These raw images were further used to crop out single cells (1000 images), budding cells (1000 images), and cell groups (1000 images) (**Figure 1**). A total of 4000 images per species consisting of raw images, single cells, budding cells, and cell groups saved in different directories and labelled with the name of the species. Test sets consisting of 200 images each of raw images, single cell, budding cell, and cell groups for each species were acquired. This test set was not included either in the training or validation process. After each model was trained, a test set was used to evaluate model performance.

**Figure 1:**
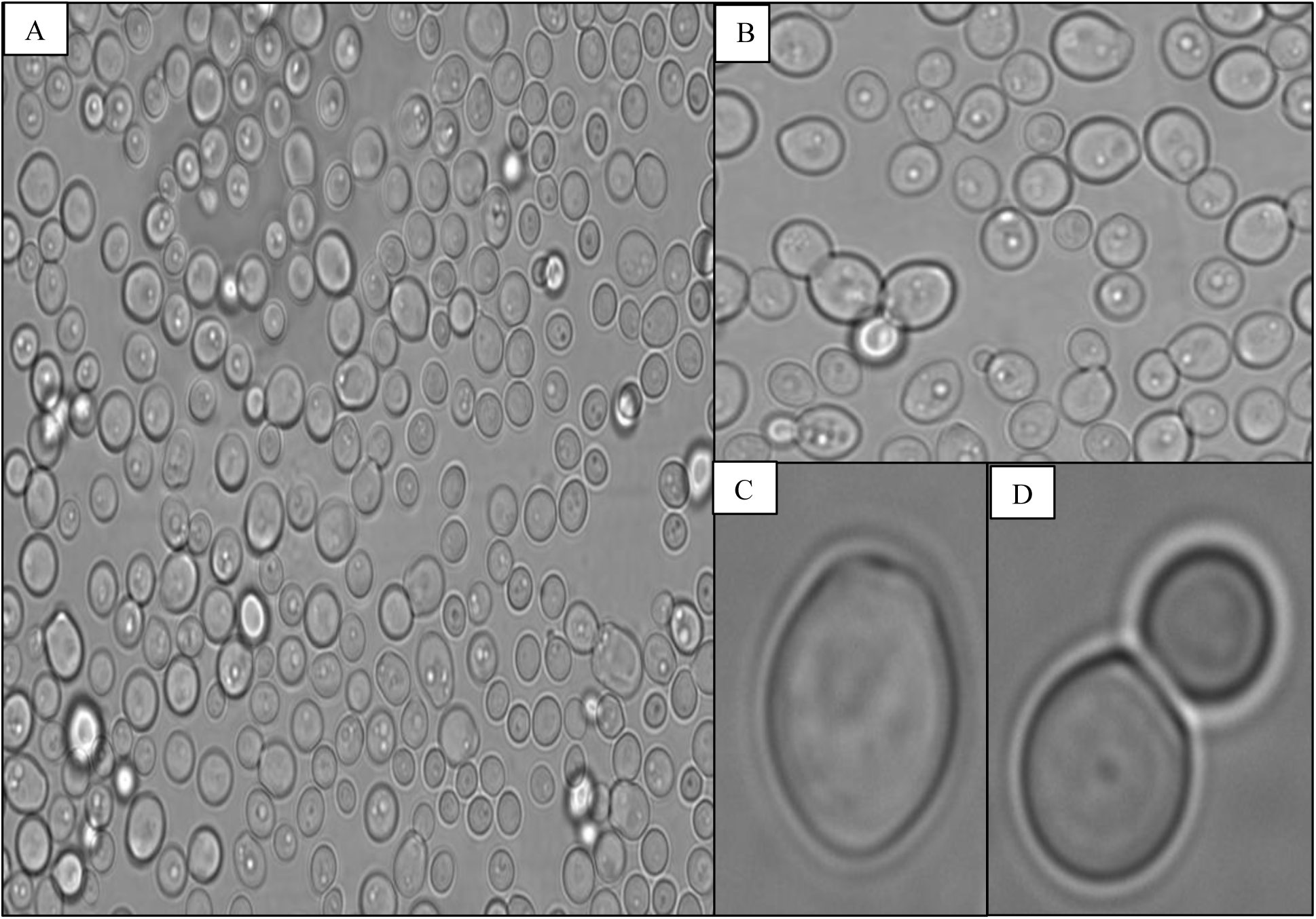
Images of *Candida* species captured using live-cell bright-field microscopy to train the machine learning models. (A) Raw microscopic image of *Candida* species visualized by wet-mount preparation after 24 hours of growth and captured using an EVOS M7000 microscope imaging system. (B) Microscopy image showing cell groups cropped from the raw image. (C) Image showing a single cell cropped from the raw image. (D) Image showing a budding cell cropped from the raw image. All these images were used in their original aspect ratio to train the deep neural network models; some of these images appear blurry here as they have been combined with others to form a montage.

### Preprocessing

All acquired microscopy images were scanned manually for corrupt or incomplete images, which were removed from the dataset. Images deemed problematic (e.g., images with scarce cells, out of focus or blurred images, etc.) to the learning task were removed. The Python Programming Language (RRID:SCR_008394) and relevant packages (Numpy, Pandas, Matplotlib, Pillow, OpenCV, TensorFlow, Keras, and Scikit-learn, scikit-image) was used for all preprocessing, simulation, and machine learning. All the images were resized to be consistent with the neural network used ^57^. Then the pixels in the images were scaled to the [0 1] range. This normalization helps to stabilize the training process and ensures that deep neural networks learn from similar intensity ranges across different images ^58^. However, for the Inception ML model images were rescaled to the pixel values from the original range [0 255] to the range of [-1 1] or [-0.5 0.5].

Mean subtraction was done to subtract the mean RGB value of the entire dataset from each image. Inception models often include mean subtraction as part of their preprocessing to center the data around zero, which aids in training convergence. Inception models have specific input size requirements (299 x 299 pixels), whereas EfficientNetB0 requires different image size (224 x 224 pixels). Images need to be resized to a fixed input size that matches the ML model’s architecture requirements. Rescaling the images helps to reduce the impact of extreme pixel values and allows the network to converge faster ^58^. Furthermore, data augmentation techniques were applied to artificially increase the diversity and quantity of training data. All the training images were subjected to data augmentation to increase the size of the dataset as well as to reduce the bias. Some of the common data augmentation techniques that we used were random rotations, translations, flips, and zooms. Data augmentation helps to reduce overfitting and improves the model’s ability to generalize to unseen data ^59^. Then the image datasets were split into separate subsets for training (80%) and validation (20%); a separate dataset was used for testing. The training dataset was used to train the model, the validation dataset for hyperparameter tuning (learning rate, batch size, number of epochs, network architecture, activation functions, weight initialization, dropout rate, fine tuning, different search strategies to explore hyperparameter space for best hyperparameters) and model selection, and the testing dataset was used to evaluate the final ML model’s performance. The overall workflow to classify *Candida* species is shown in **Figure 2**.

**Figure 2:**
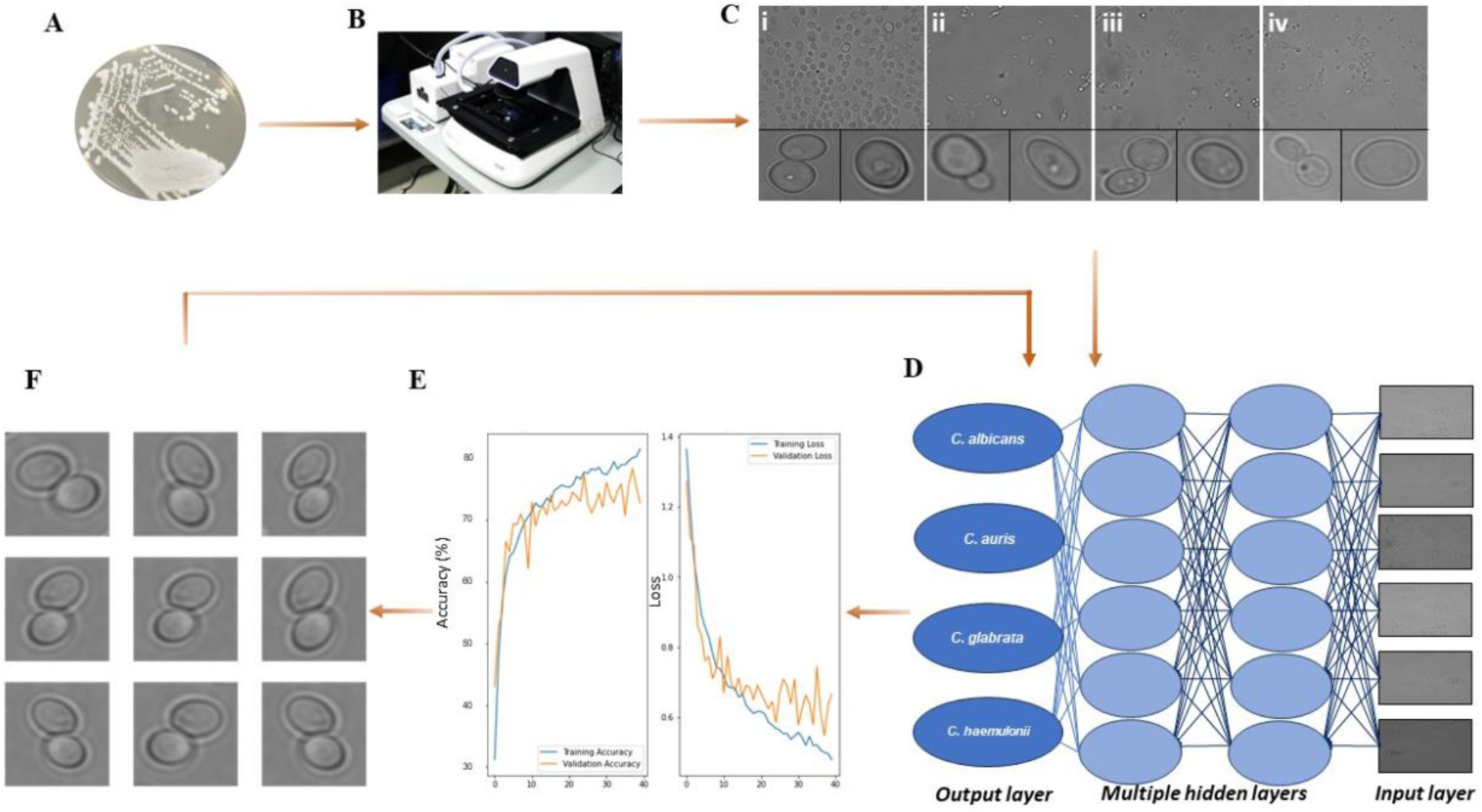
Schematic representation of the workflow to train machine learning models to identify four different *Candida* species. A) *C. albicans* on semi-solid growth medium. B) EVOS M7000 microscope imaging system. C) Labeled microscopy images of the different *Candida* species used in this study: i) *C. albicans*, ii) *C. auris*, iii) *C. glabrata* and iv) *C. haemulonii*) captured using an EVOS M7000 imaging system. D) Machine learning model reading input images and passing them through the hidden layers of the convolutional neural network to extract features and output a species classification. E) Accuracy and loss during training and validation process. F) Data augmentation (flipped, zoomed, rotated, and rescaled images of existing images) to reduce overfitting.

### Classifier Architecture

Six deep CNN classifier architectures were implemented: CNN ^60^, VGG-Net ^61^, InceptionV3 ^62^, ResNet50^63^, EfficientNetB0, and EfficientNetB7 ^64^. The custom CNN learning architectures consist of convolutional layers, pooling layers, and fully connected layers, which are stacked together in a sequential manner (**Figure S1**). There are four convolutional layers with different filters. The first convolutional layer has 16 filters, the second has 32 filters, the third has 64 filters, and the fourth has 128 filters. All these layers use a ReLU activation function. Following each convolutional layer, a max pooling layer is applied. After the last max pooling layer, a dropout layer with a dropout rate of 0.2 is added. Then the model flattens the 2D feature maps into a 1D vector. Two fully connected (dense) layers follow the flattened layer. The first dense layer consists of 128 units and uses the ReLU activation function. The second dense layer is the output layer with Softmax activation function, which has 4 class units (once for each or the *Candida* species in our study). A total of 50 epochs were used to train the model.

The VGG16 base model was created using the VGG16 class; the weights parameter was set to “ImageNet”, which initializes the model with pre-trained weights on the ImageNet dataset ^65^. Then the fully connected layers at the top of the VGG16 architecture were excluded (Figure S2). The shape of the input images was set to 224 x 224 x 3 pixels for RGB images with dimensions 224 x 224 pixels. The model was fine-tuned by freezing the pre-trained layers so that the learned features were retained. A new model was then created, and the base model added as the first layer in the new model, followed by a flattening of the layer. Two fully connected (dense) layers were added after the flattened layer. The first dense layer had 64 units and the ReLU activation function was used, which introduced non-linearity to the model. The second dense layer was the output layer, which had 4 units corresponding to each of the *Candida* species used in this study. The SoftMax activation function was used to produce class probabilities for multi-class classification. The SoftMax function ensures that the output probabilities sum up to 1.

InceptionV3, the third version of the Inception architecture (**Figure 3**) ^65^, has fewer parameters (models with less parameters are often preferred for transfer learning due to faster convergence and better adaptation to new data) compared to the VGG architecture ^66^. As for VGG16, images are scaled to 224 x 224 x 3 pixels. Images pass through each block layer then max pooling is performed at the next layer. Images pass via these convolutional blocks and into flatten layer, which is fully linked. Four dense layers (512, 256, 128, and 4 nodes) and three dropout layers (0.5) after the first three dense layers were used. The Softmax activation function was used for the output layer.

**Figure 3:**
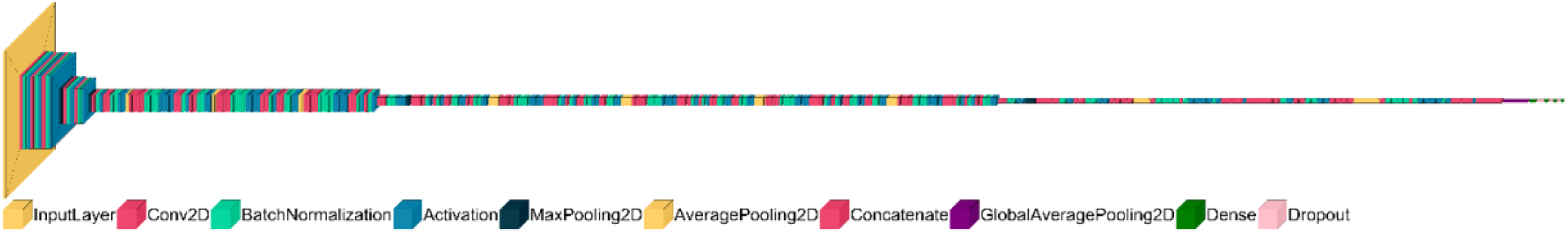
Schematic representation of InceptionV3 model. The color map defines the different layer types in the visualization (input, convolution, batch normalization, activation, max pooling, average pooling, concatenation, global average pooling, dense, and dropout layers).

ResNet50 stands out for its deep architecture, consisting of 50 layers (**Figure S3**), which enables it to learn highly complex representations from images ^63^. ResNet50 processes images of size 256 x 256 x 3 pixels. Max pooling was used to downsample the features. After passing through the residual blocks, features were flattened to a 1D vector. Like the InceptionV3 architecture, four dense layers were utilized with the final layer consisting of four nodes representing classes. Random dropout layers were also implemented after the first three layers.

EfficientNetB0 has excellent generalization capabilities across various datasets ^64^. The architecture employs a compound scaling technique that efficiently scales up the model’s dimensions while keeping computational resources in check. Whereas EfficientNetB7 represents the most advanced version of the EfficientNet CNN architecture, it has a larger model size and complexity compared to EfficientNetB0. Despite its increased computational cost, EfficientNetB7 maintains the inherent efficiency that is characteristic of the EfficientNet architecture.

## Results

We evaluated the effectiveness of the six CNN models to identify four different *Candida* species. First, we evaluated the custom CNN model with and without data augmentation, whereas all other CNN models were evaluated with data augmentation. Data augmentation artificially increases the number of images, improves model generalization, reduces overfitting and variance to transformations, and overall improves performance on real-world data. The training accuracy without data augmentation for the custom CNN model reached up to 99.6%, whereas the validation accuracy was 66.9%. Post data augmentation, the training and validation accuracy were respectively 85.4% and 83.9%, respectively (**Figure S4**). However, the precision and recall for this model were 25.0% and 38.2%, respectively. Overall, the performance of the custom CNN model on the unseen test dataset was not satisfactory (**Table S1**). The raw images contain blank spaces, which this model received as “garbage data” (i.e., data without any value towards predicting the output). Therefore, we cropped out images of single, budding, and cell groups from the raw images and then trained the model (see “Data Collection” section in supplemental materials). This model trained on the augmented data was able to identify *C. albicans* on raw cell images and budding cells respectively at 75.0% and 82.0%, though at the cost of poorly identifying *C. albicans* from single cell images (12.5%). Overall, this model performed poorly in identifying *C. haemulonii* in all the testing datasets (**Table S2**).

**Figure 4:**
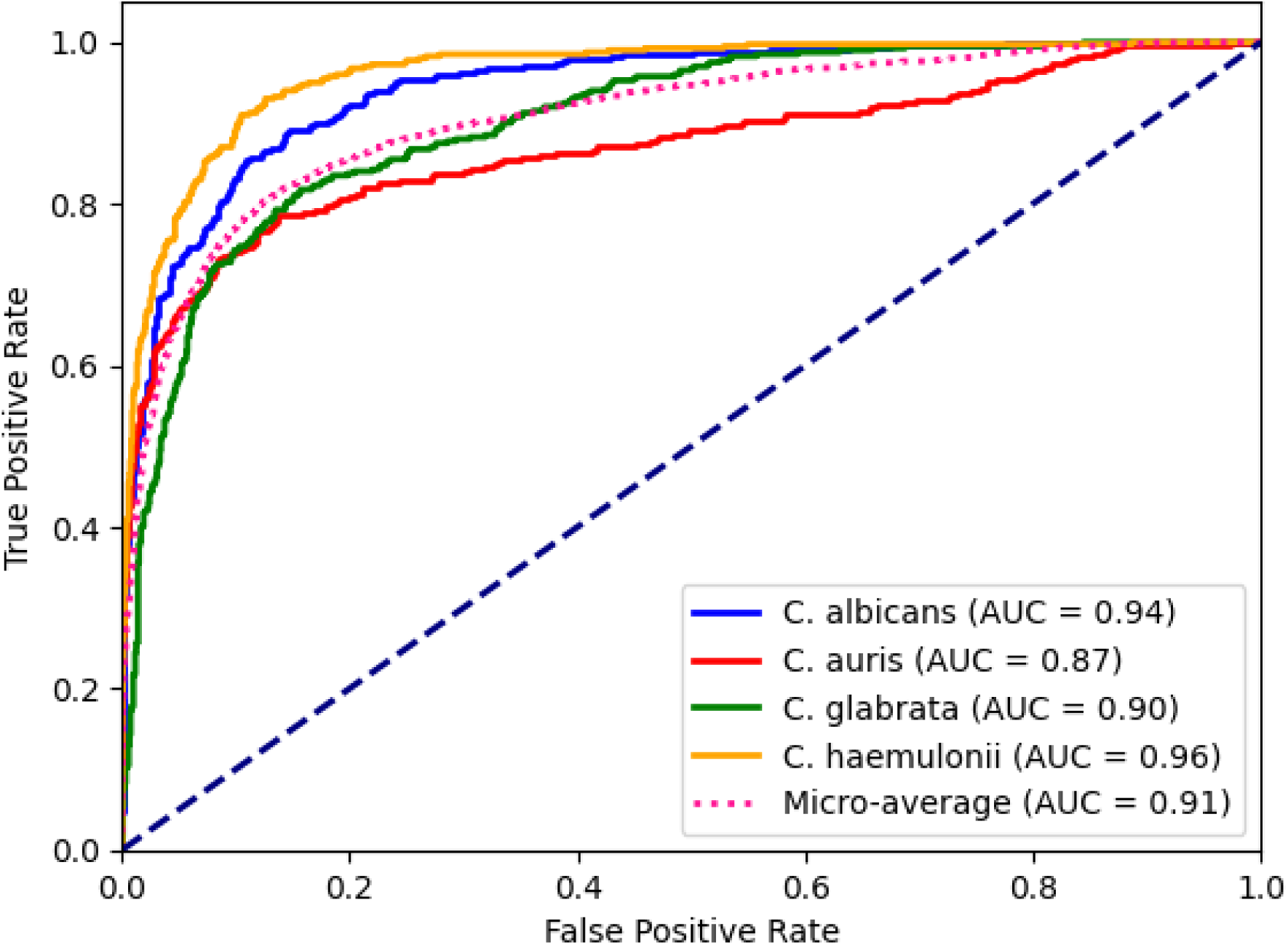
AUC-ROC curve of InceptionV3 model and AUC value with individual class scores. Dashed blue line (no-discrimination line) represents the performance of a random guessing classifier.

**Table 2:**
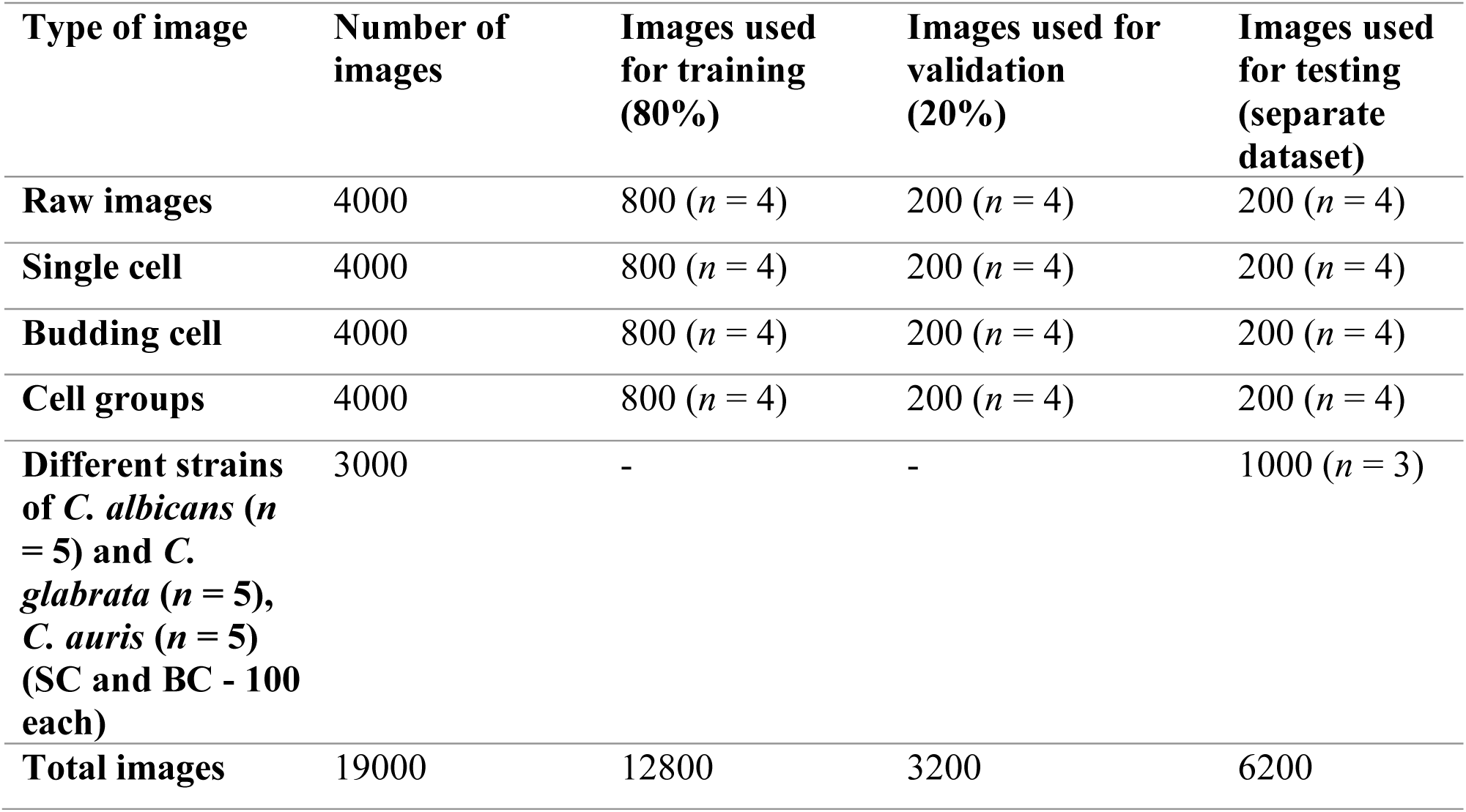
Size of the image datasets used for training, validating, and testing each model (custom CNN, InceptionV3, VGG16, ResNet50, EfficientNetB0, and EfficientNetB7) for 4 different *Candida* species. A total of 16000 images were used (1000 each of raw images, single cell images, budding cell images, and cell group images for each species). Of these 16000 images 80% (12800 images) were used for training and 20% (3200 images) were used for validation. A separate dataset consisting of 200 images for each image type (raw images, single cell images, budding cell images, and cell group images) were used for testing the models (200 x 4 (4 image types) x 4 (4 different *Candida* species) = 3200). Five different strains for each of *C. albicans*, *C. glabrata*, and *C. auris* were tested for single cell images (100 images per strain, 500 images per species) and budding cells images (100 images per strain, 500 images per species). SC denotes single cell BC denotes budding cell images.

| Type of image | Number of images | Images used for training (80%) | Images used for validation (20%) | Images used for testing (separate dataset) |
| --- | --- | --- | --- | --- |
| Raw images | 4000 | 800 ($n = 4$) | 200 ($n = 4$) | 200 ($n = 4$) |
| Single cell | 4000 | 800 ($n = 4$) | 200 ($n = 4$) | 200 ($n = 4$) |
| Budding cell | 4000 | 800 ($n = 4$) | 200 ($n = 4$) | 200 ($n = 4$) |
| Cell groups | 4000 | 800 ($n = 4$) | 200 ($n = 4$) | 200 ($n = 4$) |
| Different strains of *C. albicans* ($n = 5$) and *C. glabrata* ($n = 5$), *C. auris* ($n = 5$) (SC and BC - 100 each) | 3000 | - | - | 1000 ($n = 3$) |
| Total images | 19000 | 12800 | 3200 | 6200 |

Next, we trained the ResNet50 model with data augmentation. With data augmentation the training and validation accuracy respectively reached 98.7% and 73.0% at the end of epoch 20 (**Figure S5)**. However, the precision and recall for each class was less than 30.0% (**Table S4**).

The test set containing the raw images was used to test this model’s predictive ability. Trained ResNet50 was able to predict all the *C. albicans* (100.0%) raw images. However, it could only predict *C. haemulonii*, *C. glabrata*, and *C. auris* in 18.0%, 6.9%, and 0.5% of the cases, respectively. We then used the test set containing the single cell images. The model correctly predicted *C. albicans, C. auris, C. glabrata*, and *C. haemulonii* in 88.5%, 71.0%, 62.5%, and 40.0% instances, respectively. The model was able to predict a test set containing *C. albicans* budding cells at 96.0%, followed by *C. haemulonii* (61%), *C. glabrata* (28.4%), and *C. auris* (28.4%).

Then we trained the InceptionV3 model with data augmentation. The training and validation accuracy reached 92.4% and 78.7%, respectively. The precision and recall for this model trained and validated on all the image sets is provided in **Table S5**. When we evaluated the trained InceptionV3 model with a test set containing raw images this model was able to correctly classify the majority of *C. albicans* images (95.5%). However, this model could not identify *C. auris* and *C. haemulonii,* as the images were misclassified as *C. albicans*. A low success rate of 29.2% was also observed in correctly classifying the *C. glabrata* images. When the test set containing only budding cells was evaluated this model was able to correctly identify *C. albicans*, *C. auris*, *C. glabrata*, and *C. haemulonii* budding cells in 87.7%, 20.0%, 77.0%, and 82.3% of cases, respectively. Whereas, in the case of test set images of single cell belonging to *C. albicans*, *C. auris, C. glabrata*, and *C. haemulonii* were correctly identified in 87.0%, 59.0%, 85.0%, and 36.5% of cases, respectively. Then we trained this model using images of single cell and budding cell belonging to different classes and found that the training and validation accuracies reached 76.3% and 74.6 %, respectively, at the end of epoch 50. We then evaluated other metrics (negative predictive value, F1 score, AUC score, time to identify) for this model (**Table 3)**. The receiver operator characteristic (ROC) curve is a plot that depicts the trade-off between the sensitivity and specificity across a series of cut-off points when the diagnostic test is continuous or on ordinal scale ^67^. The ROC curve for the different *Candida* species is presented in **Figure 4**. When we tested the model’s performance on the test set containing single cells and budding cells of different *Candida* species this model could respectively identify budding cells of *C. albicans, C. auris*, *C. glabrata*, and *C. haemulonii* in 97.0%, 74.0%, 68%, and 66% of the cases. Similarly, for single cell test set model could respectively identify *C. albicans, C. auris*, *C. glabrata* and *C. haemulonii* in 97.0%, 73.0%, 69%, and 73% of the cases.

**Table 3:** Results of InceptionV3 model and for each class trained on single and budding cell images. See “Definitions” section for quantitative definitions of each metric.

| **Metrics** | **InceptionV3** | *C. albicans* | *C. auris* | *C. glabrata* | *C. haemulonii* |
| --- | --- | --- | --- | --- | --- |
| **Accuracy (%)** | 77.19 | 74.6 | 74.6 | 74.6 | 74.6 |
| **Sensitivity (%)** | 77.1 | 88.5 | 63.0 | 0.55 | 91.2 |
| **Specificity (%)** | 92.4 | 86.2 | 96.2 | 96.1 | 87.5 |
| **Precision (%)** | 77.9 | 68.2 | 84.8 | 82.8 | 70.8 |
| **Negative predictive value (%)** | 92.4 | 95.7 | 88.6 | 86.7 | 96.7 |
| **F<sub>1</sub> score (%)** | 77.14 | 77.0 | 72.3 | 66.6 | 79.7 |
| **AUC (%)** | 91.9 | 94.3 | 87.0 | 90.1 | 96.1 |
| **Time to identify (ms)** | - | 0.02 | 0.02 | 0.02 | 0.02 |

We then evaluated the VGG16 model to determine if it could outperform the InveptionV3 model. When VGG16 model was trained on all the image datasets and the overall training accuracy was 73.0% (**Table S6**). We found a similar accuracy (73.7%) when the VGG16 model trained on single cell and budding cell image dataset alone (**Table S7**). EfficientNetB0 and EfficientNetB7 models were also trained for raw images as well as single and budding cell images. However, the precision and recall for both models were below 25.0%.

Finally, we tested the ability of the InceptionV3 model to identify five different strains of the same species for *C. albicans*, *C. glabrata*, and *C. auris*. Of the total 500 images of single cells and budding cells each belonging to each strain (100 images of single cells and budding cells for each strain) the trained InceptionV3 model was able to identify accurately single cell images of *C. albicans*, *C. glabrata*, and *C. auris* in 93.4%, 69.2%, and 73.5% of the cases, respectively. Whereas for the budding cell images the model was able to identify *C. albicans*, *C. glabrata*, and *C. auris* in 92.0 %, 75.4%, and 68.3% of the cases.

## Discussion

Image based studies incorporating ML algorithms have successfully been implemented to identify disease states, quantify biomarkers, detect mitosis, recognize lymph node metastasis, tissue segmentation, prognostication, and to predict molecular expression and treatment responses ^68^. Our proof-of-concept study demonstrates that overall microscopy images obtained using wet-mounted slides can be used to identify different *Candida* species using ML models. However, our results indicate that black spaces between cell in the raw (unprocessed) images did not add to useful information with respect to cell features. In contrast, single cell and budding cell images with minimal blank spaces proved ideal. Different *Candida* species have different cell size ranges and budding pattern ^69^, which may have been critical in for our machine learning models to differentiate between the species. Our model was able to distinguish *C. albicans* more efficiently from other tested *Candida* species. One hypothesis for misclassification of *C. glabrata* and *C. haemulonii* as *C. albicans* is due to pleomorphic nature of *C. albicans* which varies in size and shape. The varying size and shape of *C. albicans* are learnt by the ML models and while classifying other *Candida* species some of these features might resemble that of *C. albicans*, which may lead to misclassification ^69^. Further studies using stained cell images to train the machine learning models could help overcome this issue.

Performance of different ML algorithms depends on the dataset used during the learning process. Similarly, our study showed that the microscopy image dataset we generated was effectively learned by InceptionV3, which was able to classify the species of *Candida* most accurately. High performance of InceptionV3 compared to other models has been reported previously. For instance, InceptionV3 showed high accuracy and sensitivity to diagnose poultry diseases ^70^.

Another study on ankle fracture detection using deep learning algorithms found that InceptionV3 model had high specificity and sensitivity ^71^. Similarly, InceptionV3 was able to identify skin cancer-melanoma with high accuracy ^72^. However, custom CNN, VGG16, ResNet50, EfficientNetB0, and EfficientNetB7 were not as efficient as InceptionV3 in classifying *Candida* species. Such similar instances of poor performance of these deep neural networks have also been found previously ^73–75^. Some of these models performed well on the other datasets such as ImageNet, CIFAR (Canadian Institute for Advanced Research) and MNIST (Modified National Institute of Standards and Technology) databases but failed to learn on the microscopy images in this study. In our study, we showed that the trained model was able to identify the images within 0.02 millisecond (**Table 3**). In contrast, diagnostic labs inoculate clinical samples onto agar plate or automated blood culture system (blood samples) and the resulting growth after incubating for 18-24 hours is utilized for the identification^76^. Identifying the etiologic agent may further take around 24 to 48 hours ^76^. Whereas the sample preparation method used in this study is comparatively easy with limited required resources after we obtain the yeast colonies on the agar plate.

One common challenge with ML models is their “black box” nature, which makes it difficult to understand how they arrive at their predictions^77^. This lack of interpretability hinders their adoption in clinical practice where transparency and trust are crucial. To address this challenge, researchers in the field of explainable AI have explored different approaches to make ML models more interpretable ^78–82^. For instance, by generating heatmaps or saliency maps, identifying informative features through feature reduction or selection, or providing immersive visualizations through VR technology, researchers can enhance transparency and trust in ML predictions. We speculate that the subtle uniqueness of each species in their shape, size, and the budding pattern along with the budding base might have contributed as the important features for ML process. Similarly, further exploration of the model’s ability to predict different *Candida* species will provide more insight into the features of the images required to identify them accurately. Based on this proof of concept study, we can further elaborate the use of these ML models in many areas such as 1) extrapolating the application of ML approach to detect other yeast species which in turn can help its diagnostic utility, 2) employing different complex models (e.g., DenseNet, Xception, MobileNetV2, ResNeXt, and SENet ^83^) to improve performance metrics, 3) location specific training of the yeast images is required for its use, and 4) differentiating antifungal resistant isolates from susceptible isolates.

One of the limitations of our study is that we did not consider all the yeast species causing infection (e.g., *Pichia kudriavzevii* (*C. krusei*), *C. parapsilosis*, and *C. tropicalis*). We also did not test microscopy images obtained from co-culturing different *Candida* species. In clinical settings multi-species infections are encountered and are challenging to treat ^84^. Distinguishing yeast species from other cell types and components of clinical samples such as blood and urine would further elaborate the utility of our method for clinical applications. Nevertheless, we anticipate that our machine learning-based approach, which does not require sophisticated instruments or extensive expertise to implement, will help to reduce the turnaround times for the diagnosis of yeast infections.

Our study demonstrates that the identification time for *Candida* species can be drastically reduced and requires less expertise unlike for sophisticated techniques such as MALDI-TOF MS or nucleic acid-based amplification tests. Our study also highlights the ease of use in exploiting ML approach for rapid identification of *Candida* species using microscopy images. Considering the rapid emergence of antifungal-resistant species ^33^, it will be indispensable to evaluate the ability of these ML models to determine susceptibility profiles in future work. For instance, data from the whole genome sequence of *C. auris* has been used to train ML models to predict the mutations responsible for antifungal drug resistance, rank different resistance mutations, and discover new mutations related to drug resistance ^85^. Overall, we anticipate that ML approaches will be able to detect and determine antifungal-resistant phenotypes, underlying molecular resistance mechanisms, and improve the treatment of patients with infectious fungal diseases.

## Author contributions

Conceptualization, D.A.C.; methodology, S.A.S. and D.A.C.; experimental investigation, S.A.S.; formal analysis and visualization, S.A.S.; data curation, S.A.S.; supervision, D.A.C.; funding acquisition, D.A.C.; S.A.S. and D.A.C. wrote and approved the published version of the manuscript.

## Acknowledgements

We thank Dr. Tanis Dingle and the Alberta Precision Laboratories-Public Health Laboratory for providing the *C. auris* isolates. We acknowledge the Samira Rasouli Koohi and the Molecular Biology Services Unit at the University of Alberta for assistance with genetic sequencing. We acknowledge Prof. Randy Goebel, Prof. Jay Newby, and Jiyai Dai for guidance on choosing, training, and validating the machine learning models. We thank Joshua Guthrie for technical support and proofreading the manuscript.

## Data Sharing and Data Accessibility

The data that support the findings of this study are available upon reasonable request from the corresponding author.

## Supplementary Materials

### Definitions

*Specificity* ^86^: Specificity, also known as the “true negative rate”, measures the proportion of true negative predictions among all actual negative instances in the dataset. It indicates how well a model can correctly identify the negative cases. Specificity is calculated using the formula:

Specificity = True Negatives / (True Negatives + False Positives)

*Sensitivity* (Recall or True Positive Rate)^86^: Sensitivity measures the proportion of true positive predictions among all actual positive instances in the dataset. It indicates how well a model can identify all the positive cases. Sensitivity is calculated using the formula:

Sensitivity = True Positives / (True Positives + False Negatives)

*Precision* ^87^: Precision is a metric that measures the proportion of true positive predictions among all positive predictions made by a model. It tells how many of the items predicted as positive are true positives. Precision is calculated using the formula:

Precision = True Positives / (True Positives + False Positives)

*Recall* (Sensitivity or True Positive Rate) ^87^: Recall is a metric that measures the proportion of true positive predictions among all actual positive instances in the dataset. It indicates how well a model can identify all the positive cases. Recall is calculated using the formula:

Recall = True Positives / (True Positives + False Negatives)

*F1 Score* ^88^: The F1 score is a harmonic mean (the reciprocal of the arithmetic mean of the reciprocals) of precision and recall. It provides a balanced measure that considers false positives and false negatives. The F1 score is useful when you want to find a balance between precision and recall. It is calculated using the formula:

F1 Score = 2 * (Precision * Recall) / (Precision + Recall)

*Area Under the ROC Curve* (AUC) ^89^: The ROC curve (receiver operating characteristic curve) is a graphical representation of the performance of a classification model at different classification thresholds. AUC measures the area under this curve and provides an aggregated measure of a model’s ability to discriminate between positive and negative instances. AUC values range between 0 and 1, with higher values indicating better performance.

*Negative Predictive Value* (NPV) ^88^: The Negative Predictive Value is a metric that represents the proportion of true negative predictions among all negative predictions made by a model.

NPV is useful for understanding a model’s performance in identifying true negatives. It is calculated using the formula:

NPV = True Negatives / (True Negatives + False Negatives)

## Supplementary Figures

**Figure S1:**
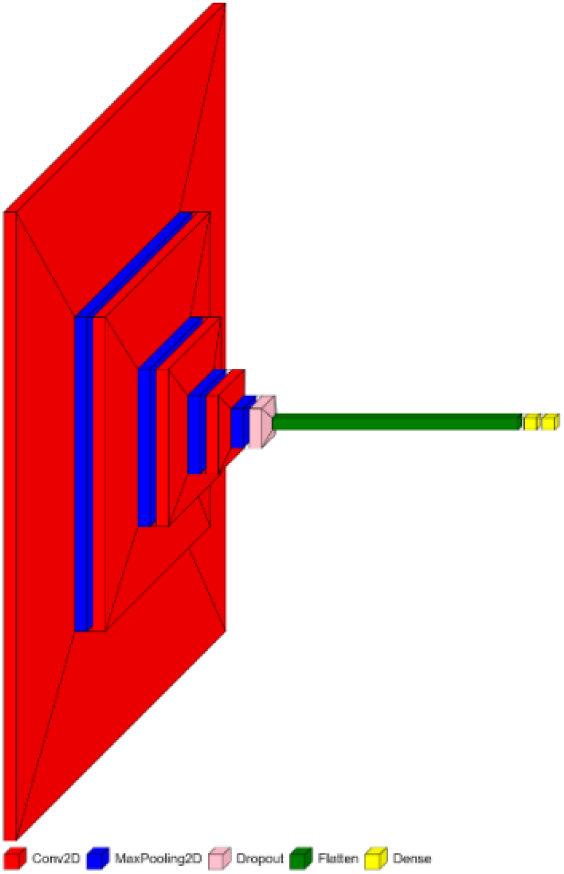
Schematic representation for visualization of the custom CNN Model. The color map defines the different layer types in the visualization (convolution, max pooling, dropout, flatten, and dense layers).

**Figure S2:**
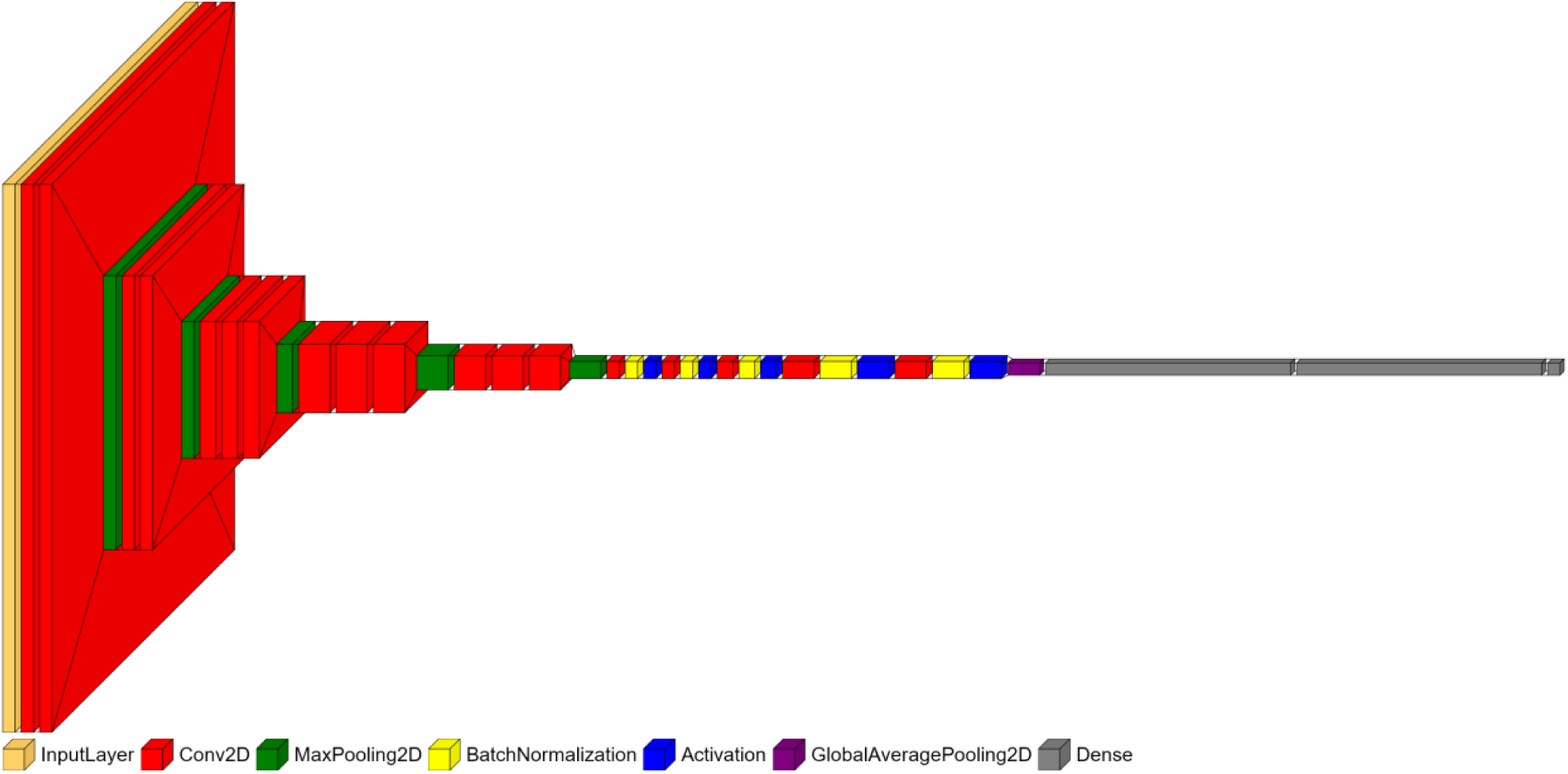
Schematic representation for visualization of the VGG16 model. The color map defines the different layer types in the visualization (input, convolution, max pooling, batch normalization, activation, global average pooling, and dense layers).

**Figure S3:**
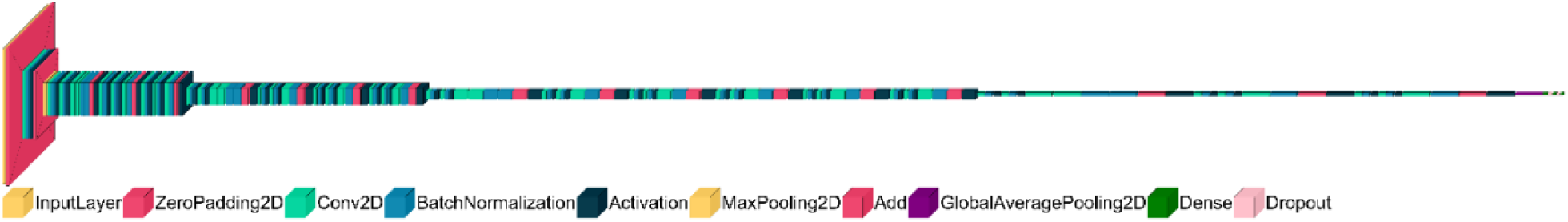
Schematic representation for visualization of the ResNet50 model. The color map defines the different layer types in the visualization (input, zeropadding, convolutional, batch normalization, activation, max pooling, adding, global average pooling, dense, and dropout layers).

**Figure S4:**
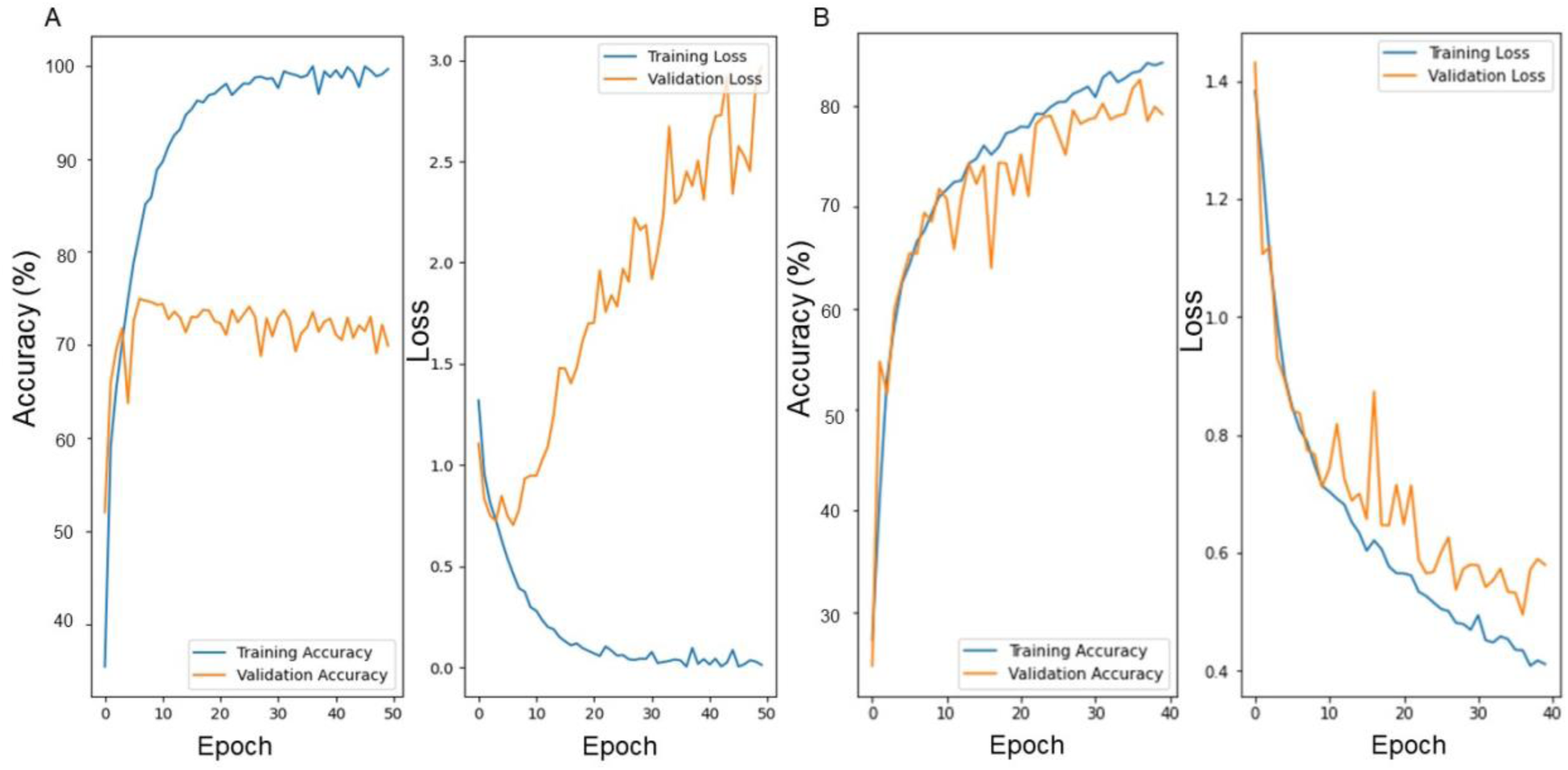
Training and validation accuracies of the custom CNN model before and after data augmentation. A) Training and validation accuracies and loss before data augmentation. B) Training and validation accuracies and loss after data augmentation.

**Figure S5:**
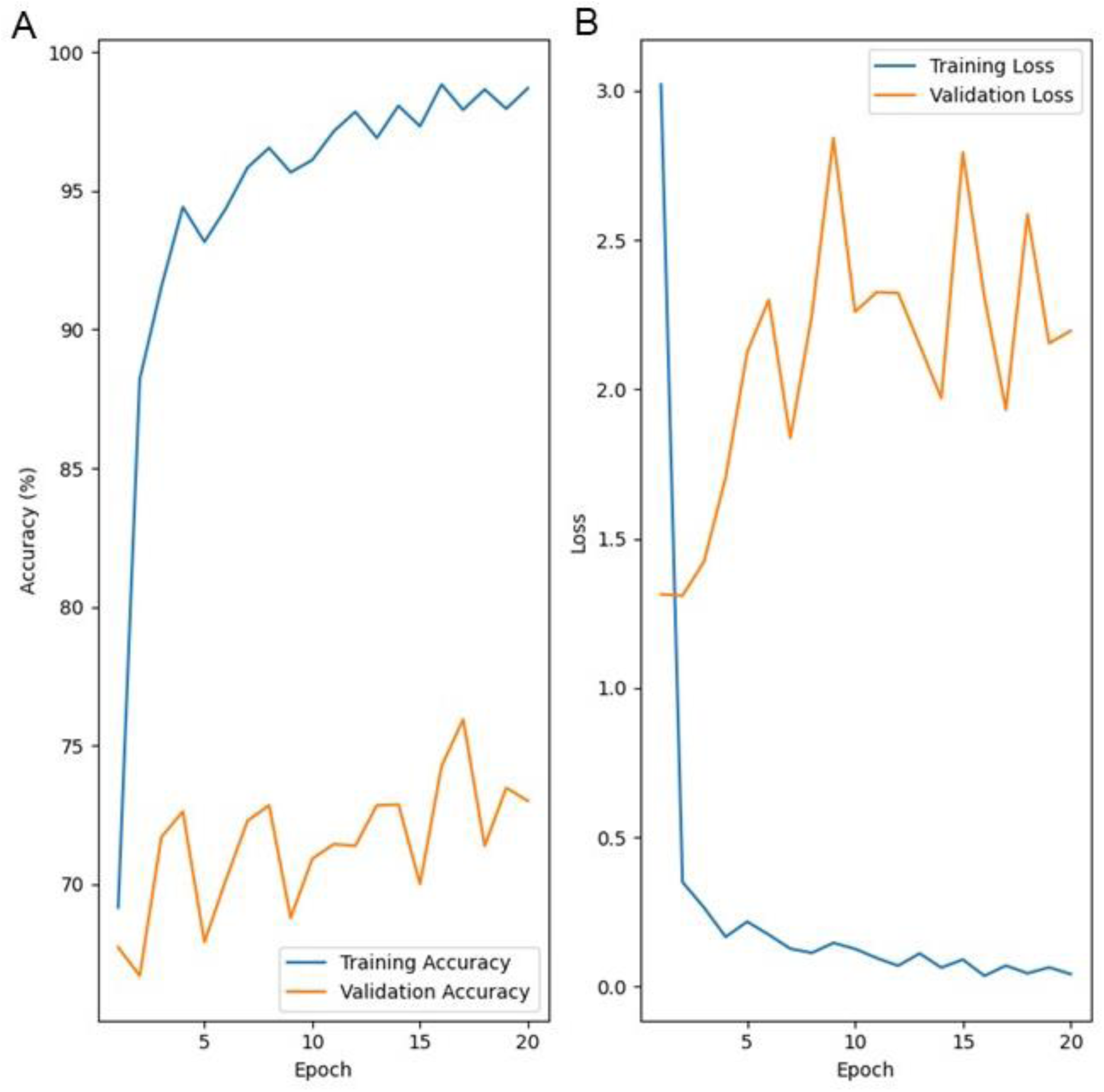
Training and validation accuracies and loss of the ResNet50 model. These metrics were evaluated for A) raw images, single cell, budding cell, and cell groups and for B) single cell and budding cell images.

**Figure S6:**
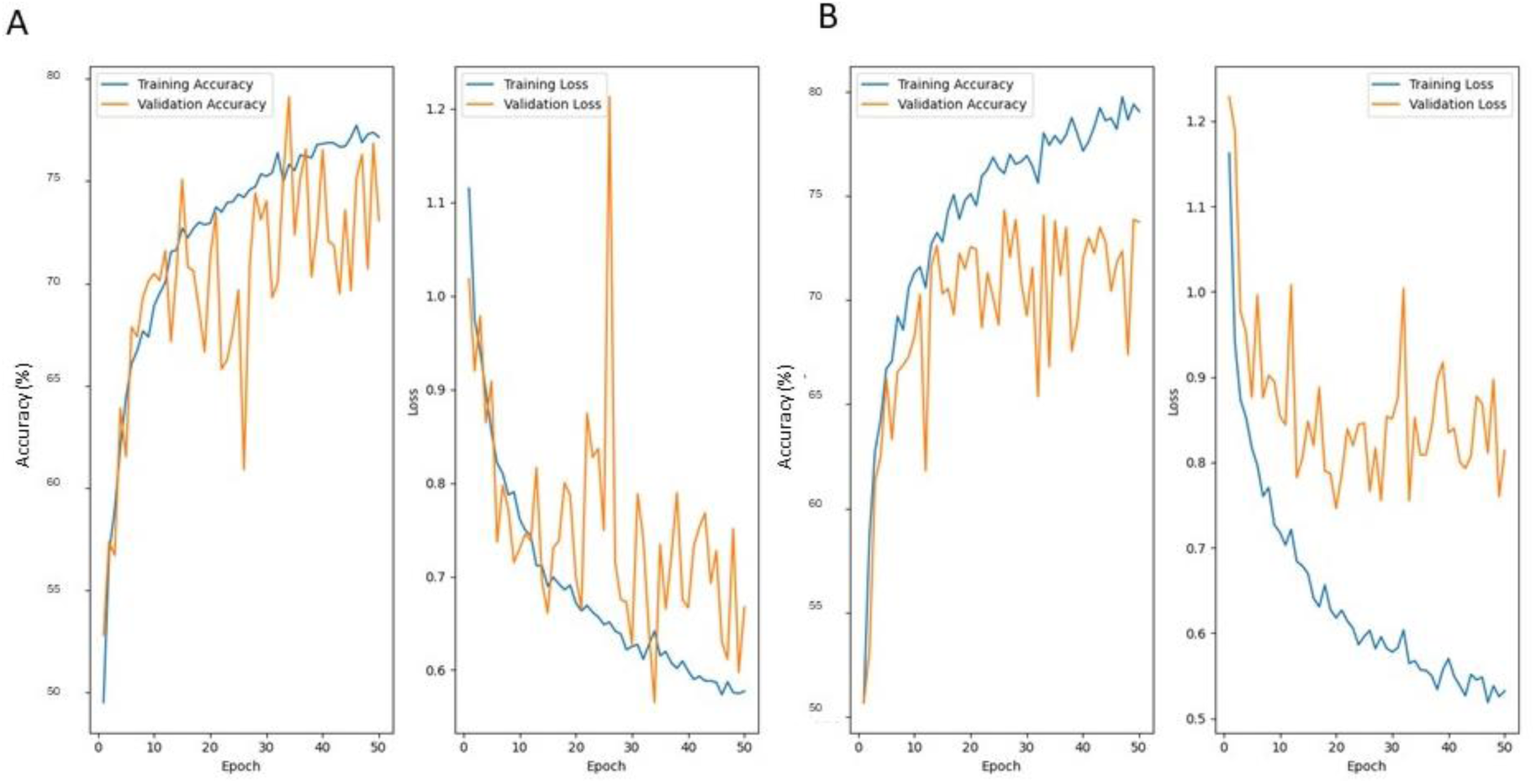
Training and validation accuracies and loss of the VGG16 model. These metrics were evaluated for A) raw images, single cell, budding cell, and cell groups and for B) single cell and budding cell images.

## Supplementary Tables

**Table S1:** Metrics of the custom CNN after data augmentation on raw cell, single cell, budding cell, and cell groups images. See “Definitions” section for quantitative definitions of each metric.

| **Metrics** | **Custom CNN**<br>**(All images)** | *C. albicans* | *C. auris* | *C. glabrata* | *C. haemulonii* |
| --- | --- | --- | --- | --- | --- |
| **Accuracy (%)** | 69.7 | 81.7 | 74.6 | 71.1 | 51.3 |
| **Sensitivity (%)** | 69.7 | 81.7 | 55.3 | 71.1 | 51.3 |
| **Specificity (%)** | 69.7 | 93.8 | 90.4 | 90.4 | 51.3 |
| **Precision (%)** | 72.0 | 78.2 | 55.3 | 72.5 | 82.0 |
| **Negative**<br>**predictive value**<br>**(%)** | 95.4 | 92.4 | 79.9 | 91.0 | 96.2 |
| **F1 score (%)** | 69.6 | 79.9 | 63.5 | 71.8 | 63.1 |
| **AUC (%)** | 87.1 | 91.1 | 81.7 | 88.3 | 87.5 |

**Table S2:** Metrics of the custom CNN after data augmentation on single cell (SC) and budding cell (BC) images. See “Definitions” section for quantitative definitions of each metric.

| **Metrics** | **Custom CNN**<br>**(SC and BC**<br>**dataset)** | *C. albicans* | *C. auris* | *C. glabrata* | *C. haemulonii* |
| --- | --- | --- | --- | --- | --- |
| **Accuracy (%)** | 69.7 | 81.7 | 74.6 | 71.1 | 51.3 |
| **Sensitivity (%)** | 69.7 | 81.7 | 74.6 | 71.1 | 51.3 |
| **Specificity (%)** | 85.5 | 93.8 | 90.4 | 90.1 | 85.5 |
| **Precision (%)** | 72.0 | 78.2 | 55.3 | 72.5 | 82.0 |
| **Negative**<br>**predictive value**<br>**(%)** | 94.3 | 92.4 | 79.9 | 91.0 | 96.2 |
| **F<sub>1</sub> score (%)** | 69.6 | 79.9 | 63.5 | 71.8 | 63.1 |
| **AUC (%)** | 87.1 | 81.7 | 81.7 | 88.3 | 87.5 |

**Table S3:**
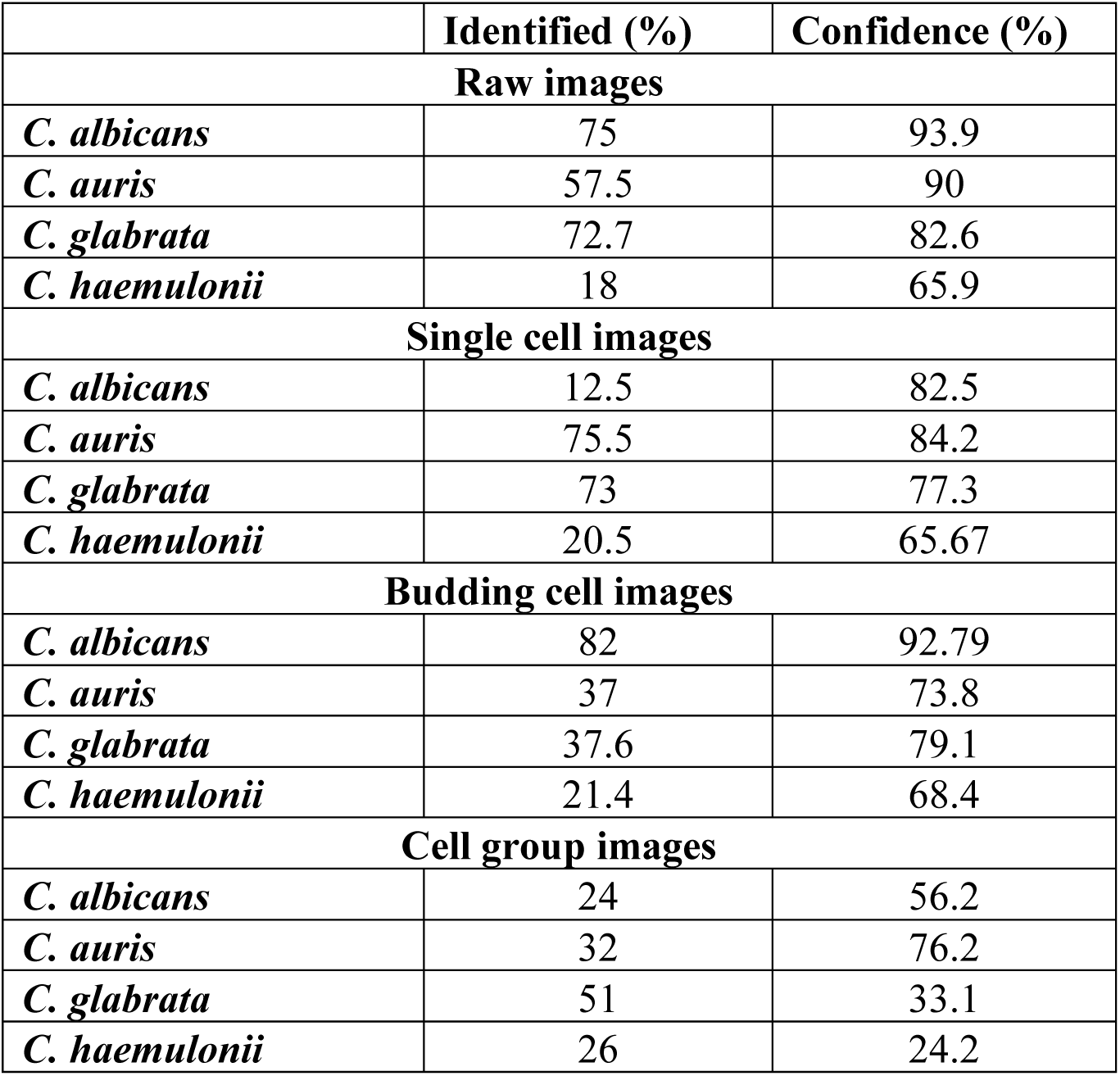
Performance of the custom CNN model on different test image sets of four *Candida* species.

**Table S4:**
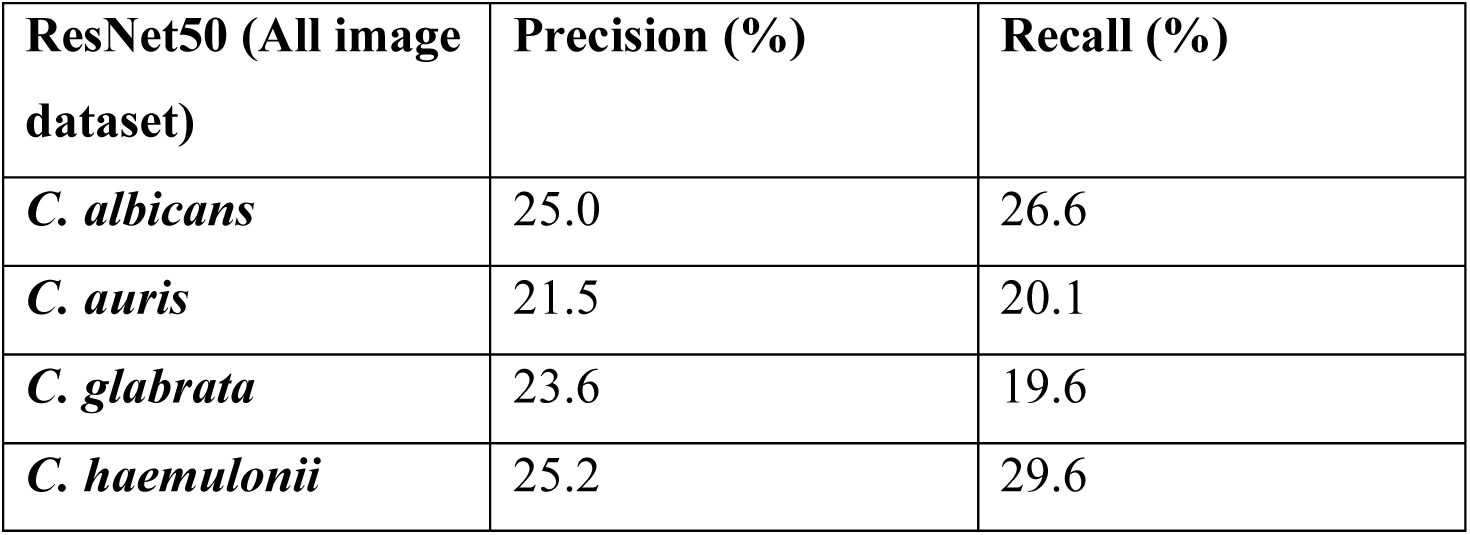
Precision and recall values for the four yeast species on the validation set for the trained ResNet50 model.

**Table S5:** Precision and recall for four different *Candida* species after training InceptionV3 for all images and training InceptionV3 on single and budding cell images together.

|  | With data augmentation (single and budding cell datasets) |  |
| --- | --- | --- |
|  | Precision (%) | Recall (%) |
| *C. albicans* | 68.2 | 88.5 |
| *C. auris* | 84.8 | 63.0 |
| *C. glabrata* | 82.8 | 55.7 |
| *C. haemulonii* | 70.8 | 91.2 |

**Table S6:** Results of VGG16 model and for each class of *Candida* species trained on raw cells, single cell, budding cell, and cell groups images. See “Definitions” section for quantitative definitions of each metric.

| Metrics | VGG16 | *C. albicans* | *C. auris* | *C. glabrata* | *C. haemulonii* |
| --- | --- | --- | --- | --- | --- |
| Accuracy (%) | 73.0 | 93.3 | 84.2 | 51.8 | 61.0 |
| Sensitivity (%) | 72.6 | 93.3 | 84.2 | 51.8 | 61.0 |
| Specificity (%) | 72.8 | 80.0 | 72.1 | 77.6 | 61.5 |
| Precision (%) | 73.1 | 80.0 | 72.1 | 77.6 | 61.5 |
| Negative predictive value (%) | 95.1 | 93.3 | 84.2 | 51.8 | 61.0 |
| F1 score (%) | 72.2 | 86.1 | 77.6 | 62.1 | 61.3 |
| AUC (%) | 92.7 | 98.0 | 94.6 | 90.0 | 88.0 |

**Table S7:**
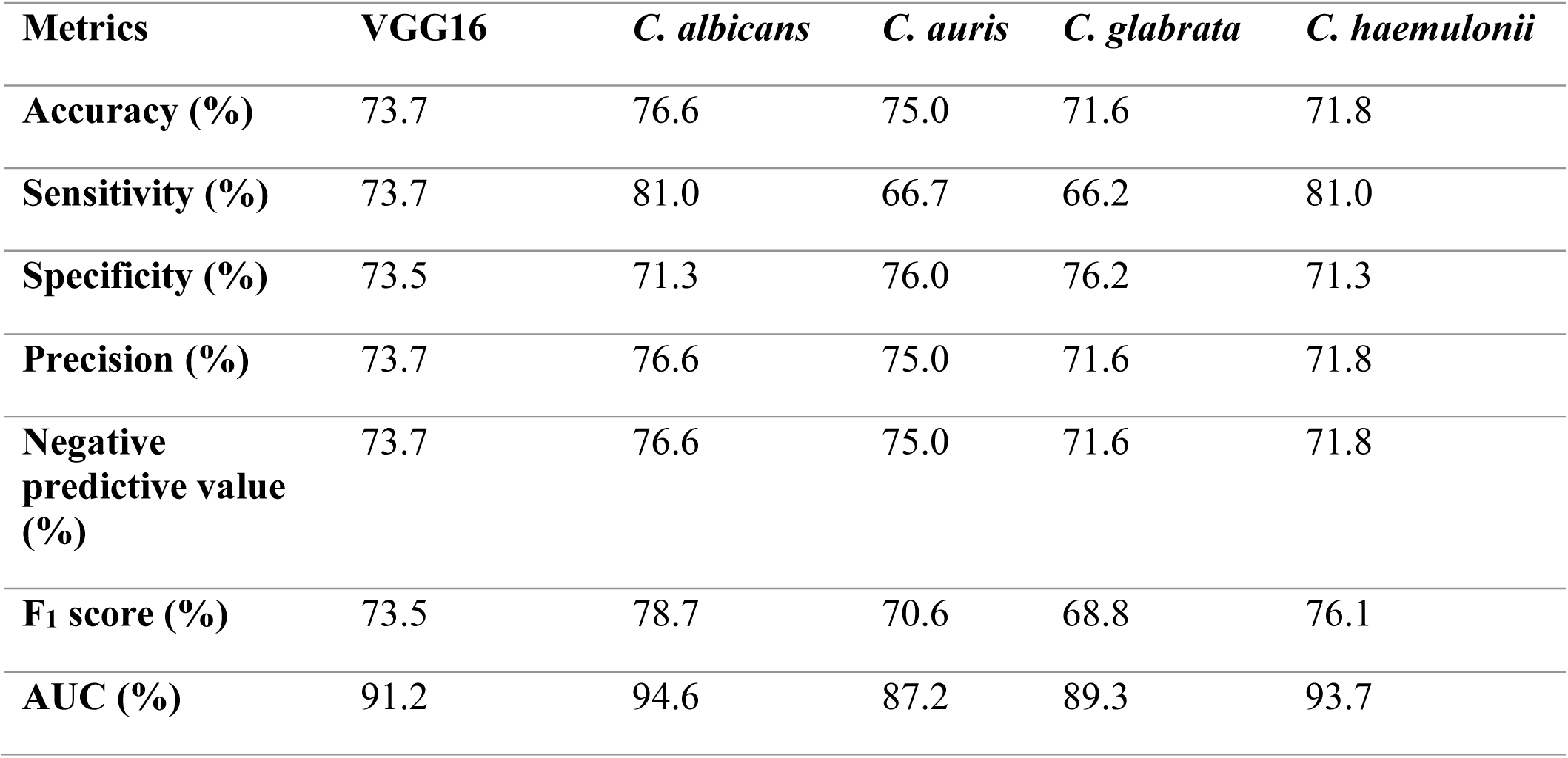
Results of VGG16 model and for each class trained on single cell and budding cell images alone. See “Definitions” section for quantitative definitions of each metric.

| **Metrics** | **VGG16** | ***C. albicans*** | ***C. auris*** | ***C. glabrata*** | ***C. haemulonii*** |
| --- | --- | --- | --- | --- | --- |
| **Accuracy (%)** | 73.7 | 76.6 | 75.0 | 71.6 | 71.8 |
| **Sensitivity (%)** | 73.7 | 81.0 | 66.7 | 66.2 | 81.0 |
| **Specificity (%)** | 73.5 | 71.3 | 76.0 | 76.2 | 71.3 |
| **Precision (%)** | 73.7 | 76.6 | 75.0 | 71.6 | 71.8 |
| **Negative predictive value (%)** | 73.7 | 76.6 | 75.0 | 71.6 | 71.8 |
| **F1 score (%)** | 73.5 | 78.7 | 70.6 | 68.8 | 76.1 |
| **AUC (%)** | 91.2 | 94.6 | 87.2 | 89.3 | 93.7 |

## References

1. Almeida F, Rodrigues ML, Coelho C. The Still Underestimated Problem of Fungal Diseases Worldwide. Front Microbiol. 2019;10:214. doi:10.3389/fmicb.2019.00214

2. Wisplinghoff H, Bischoff T, Tallent SM, Seifert H, Wenzel RP, Edmond MB. Nosocomial bloodstream infections in US hospitals: analysis of 24,179 cases from a prospective nationwide surveillance study. Clin Infect Dis. 2004;39(3):309–317. doi:10.1086/421946

3. Cuervo G, Garcia-Vidal C, Puig-Asensio M, et al. Usefulness of guideline recommendations for prognosis in patients with candidemia. Med Mycol. 2019;57(6):659–667. doi:10.1093/mmy/myy118

4. Arendrup MC. Epidemiology of invasive candidiasis. Curr Opin Crit Care. 2010;16(5):445–452. doi:10.1097/MCC.0b013e32833e84d2

5. Romo JA, Kumamoto CA. On Commensalism of Candida. Journal of Fungi. 2020;6(1):16. doi:10.3390/jof6010016

6. Sharma C, Kadosh D. Post-transcriptional control of antifungal resistance in human fungal pathogens. Crit Rev Microbiol. Published online 2022. doi:10.1080/1040841X.2022.2080527

7. Liu F, Zhong L, Zhou F, et al. Clinical Features, Strain Distribution, Antifungal Resistance and Prognosis of Patients with Non-albicans Candidemia: A Retrospective Observational Study. Infect Drug Resist. 2021;Volume 14:3233–3246. doi:10.2147/IDR.S323583

8. Borman AM, Johnson EM. Name changes for fungi of medical importance, 2018 to 2019. J Clin Microbiol. 2021;59(2). doi:10.1128/JCM.01811-20

9. Turner SA, Butler G. The Candida pathogenic species complex. Cold Spring Harb Perspect Med. 2014;4(9). doi:10.1101/cshperspect.a019778

10. Perfect JR. The antifungal pipeline: a reality check. Nat Rev Drug Discov. 2017;16(9):603–616. doi:10.1038/NRD.2017.46

11. Du H, Bing J, Hu T, Ennis CL, Nobile CJ, Huang G. Candida auris: Epidemiology, biology, antifungal resistance, and virulence. PLoS Pathog. 2020;16(10):e1008921. doi:10.1371/journal.ppat.1008921

12. Parums D V. Editorial: The World Health Organization (WHO) Fungal Priority Pathogens List in Response to Emerging Fungal Pathogens During the COVID-19 Pandemic. Medical Science Monitor. 2022;28. doi:10.12659/MSM.939088

13. Satoh K, Makimura K, Hasumi Y, Nishiyama Y, Uchida K, Yamaguchi H. Candida aurissp. nov., a novel ascomycetous yeast isolated from the external ear canal of an inpatient in a Japanese hospital. Microbiol Immunol. 2009;53(1):41–44. doi:10.1111/j.1348-0421.2008.00083.x

14. Rudramurthy SM, Chakrabarti A, Paul RA, et al. Candida auris candidaemia in Indian ICUs: Analysis of risk factors. Journal of Antimicrobial Chemotherapy. 2017;72(6). doi:10.1093/jac/dkx034

15. Cortegiani A, Misseri G, Fasciana T, Giammanco A, Giarratano A, Chowdhary A. Epidemiology, clinical characteristics, resistance, and treatment of infections by Candida auris. J Intensive Care. 2018;6(1):69. doi:10.1186/s40560-018-0342-4

16. Shastri PS, Shankarnarayan SA, Oberoi J, Rudramurthy SM, Wattal C, Chakrabarti A. Candida auris candidaemia in an intensive care unit – Prospective observational study to evaluate epidemiology, risk factors, and outcome. J Crit Care. 2020;57:42–48. doi:10.1016/j.jcrc.2020.01.004

17. Arendrup MC, Patterson TF. Multidrug-Resistant Candida: Epidemiology, Molecular Mechanisms, and Treatment. J Infect Dis. 2017;216(suppl_3):S445-S451. doi:10.1093/infdis/jix131

18. Chowdhary A, Sharma C, Duggal S, et al. New Clonal Strain of *Candida auris*, Delhi, India. Emerg Infect Dis. 2013;19(10):1670–1673. doi:10.3201/eid1910.130393

19. Ceyssens PJ, Soetaert K, Timke M, et al. Matrix-Assisted Laser Desorption Ionization–Time of Flight Mass Spectrometry for Combined Species Identification and Drug Sensitivity Testing in Mycobacteria. J Clin Microbiol. 2017;55(2):624–634. doi:10.1128/JCM.02089-16

20. Spivak ES, Hanson KE. Candida auris: an Emerging Fungal Pathogen. J Clin Microbiol. 2018;56(2). doi:10.1128/JCM.01588-17

21. Earnshaw SR, McDade C, Bryan A, et al. Real-World Financial and Clinical Impact of Diagnostic-Driven and Empirical-Treatment Strategies in High-Risk Immunocompromised Patients with Suspected Aspergillus Infection in the United Kingdom. Microbiol Spectr. 2022;10(3). doi:10.1128/spectrum.00425-22

22. Maaroufi Y, De Bruyne JM, Duchateau V, Georgala A, Crokaert F. Early Detection and Identification of Commonly Encountered Candida Species from Simulated Blood Cultures by Using a Real-Time PCR-Based Assay. The Journal of Molecular Diagnostics. 2004;6(2):108–114. doi:10.1016/S1525-1578(10)60498-9

23. Cherkaoui A, Renzi G, Azam N, Schorderet D, Vuilleumier N, Schrenzel J. Rapid identification by MALDI-TOF/MS and antimicrobial disk diffusion susceptibility testing for positive blood cultures after a short incubation on the WASPLab. European Journal of Clinical Microbiology & Infectious Diseases. 2020;39(6):1063–1070. doi:10.1007/s10096-020-03817-8

24. Kal Çakmaklıoğulları E, Aşgın N, Değerli K. [A comparison of the costs, reliability and time of result periods of widely used methods, new molecular methods and MALDI TOF-MS in the routine diagnosis of Candida strains]. Mikrobiyol Bul. 2019;53(2):204–212. doi:10.5578/mb.67952

25. Clancy CJ, Nguyen MH. Finding the “Missing 50%” of Invasive Candidiasis: How Nonculture Diagnostics Will Improve Understanding of Disease Spectrum and Transform Patient Care. Clinical Infectious Diseases. 2013;56(9):1284–1292. doi:10.1093/cid/cit006

26. Fernández-Manteca MG, Ocampo-Sosa AA, Ruiz de Alegría-Puig C, et al. Automatic classification of Candida species using Raman spectroscopy and machine learning. Spectrochim Acta A Mol Biomol Spectrosc. 2023;290:122270. doi:10.1016/j.saa.2022.122270

27. Donnelly JP, Chen SC, Kauffman CA, et al. Revision and Update of the Consensus Definitions of Invasive Fungal Disease From the European Organization for Research and Treatment of Cancer and the Mycoses Study Group Education and Research Consortium. Clinical Infectious Diseases. 2020;71(6):1367–1376. doi:10.1093/cid/ciz1008

28. Dupuis C, Le bihan C, Maubon D, et al. Performance of Repeated Measures of (1–3)-β-D-Glucan, Mannan Antigen, and Antimannan Antibodies for the Diagnosis of Invasive Candidiasis in ICU Patients: A Preplanned Ancillary Analysis of the EMPIRICUS Randomized Clinical Trial. Open Forum Infect Dis. 2021;8(3). doi:10.1093/ofid/ofab080

29. León C, Ruiz-Santana S, Saavedra P, et al. Contribution of Candida biomarkers and DNA detection for the diagnosis of invasive candidiasis in ICU patients with severe abdominal conditions. Crit Care. 2016;20(1):149. doi:10.1186/s13054-016-1324-3

30. Chang SS, Hsieh WH, Liu TS, et al. Multiplex PCR System for Rapid Detection of Pathogens in Patients with Presumed Sepsis – A Systemic Review and Meta-Analysis. PLoS One. 2013;8(5):e62323. doi:10.1371/journal.pone.0062323

31. White PL, Hibbitts SJ, Perry MD, et al. Evaluation of a Commercially Developed Semiautomated PCR–Surface-Enhanced Raman Scattering Assay for Diagnosis of Invasive Fungal Disease. J Clin Microbiol. 2014;52(10):3536–3543. doi:10.1128/JCM.01135-14

32. Bongomin F, Gago S, Oladele RO, Denning DW. Global and Multi-National Prevalence of Fungal Diseases-Estimate Precision. J Fungi (Basel). 2017;3(4). doi:10.3390/jof3040057

33. Bhattacharya S, Sae-Tia S, Fries BC. Candidiasis and Mechanisms of Antifungal Resistance. Antibiotics. 2020;9(6):312. doi:10.3390/antibiotics9060312

34. Clancy CJ, Nguyen MH. Diagnosing Invasive Candidiasis. J Clin Microbiol. 2018;56(5). doi:10.1128/JCM.01909-17

35. Doi K. Computer-aided diagnosis in medical imaging: historical review, current status and future potential. Comput Med Imaging Graph. 2007;31(4-5):198–211. doi:10.1016/j.compmedimag.2007.02.002

36. Medical Image Analysis and Informatics: Computer-Aided Diagnosis and Therapy - Google Books. Accessed August 9, 2023. https://books.google.ca/books?hl=en&lr=&id=QGpQDwAAQBAJ&oi=fnd&pg=PP1&ots=KtPQIOlLpE&sig=iaamY2AyvLObBzdhkqxvVKl44es&redir_esc=y#v=onepage&q&f=false

37. Liu X, Faes L, Kale AU, et al. A comparison of deep learning performance against health-care professionals in detecting diseases from medical imaging: a systematic review and meta-analysis. Lancet Digit Health. 2019;1(6):e271–e297. doi:10.1016/S2589-7500(19)30123-2

38. Ahuja AS. The impact of artificial intelligence in medicine on the future role of the physician. PeerJ. 2019;7:e7702. doi:10.7717/peerj.7702

39. Zeiler MD, Fergus R. Visualizing and Understanding Convolutional Networks arXiv:1311.2901v3 [cs.CV] 28 Nov 2013. Computer Vision–ECCV 2014. 2014;8689(PART 1):818–833. doi:10.1007/978-3-319-10590-1_53

40. Krizhevsky A, Sutskever I, Hinton GE. ImageNet classification with deep convolutional neural networks. Commun ACM. 2017;60(6):84–90. doi:10.1145/3065386

41. Brownstein JS, Freifeld CC, Madoff LC. Influenza A (H1N1) Virus, 2009 — Online Monitoring. New England Journal of Medicine. 2009;360(21):2156–2156. doi:10.1056/NEJMp0904012

42. Maharaj AS, Parker J, Hopkins JP, et al. The effect of seasonal respiratory virus transmission on syndromic surveillance for COVID-19 in Ontario, Canada. Lancet Infect Dis. 2021;21(5):593–594. doi:10.1016/S1473-3099(21)00151-1

43. Pascucci M, Royer G, Adamek J, et al. AI-based mobile application to fight antibiotic resistance. Nat Commun. 2021;12(1):1173. doi:10.1038/s41467-021-21187-3

44. Sundermann AJ, Chen J, Kumar P, et al. Whole-Genome Sequencing Surveillance and Machine Learning of the Electronic Health Record for Enhanced Healthcare Outbreak Detection. Clin Infect Dis. 2022;75(3):476–482. doi:10.1093/cid/ciab946

45. Wu J, Xie X, Yang L, et al. Mobile health technology combats COVID-19 in China. J Infect. 2021;82(1):159–198. doi:10.1016/j.jinf.2020.07.024

46. Rebrosova K, Samek O, Kizovsky M, Bernatova S, Hola V, Ruzicka F. Raman Spectroscopy—A Novel Method for Identification and Characterization of Microbes on a Single-Cell Level in Clinical Settings. Front Cell Infect Microbiol. 2022;12. doi:10.3389/fcimb.2022.866463

47. Zawadzki P, Adamczuk P, Jamka K, Wróblewska-Łuczka P, Bojar H, Raszewski G. The Microorganism Detection System (SDM) for microbiological control of cosmetic products. Annals of Agricultural and Environmental Medicine. 2021;28(4):705–708. doi:10.26444/aaem/144668

48. Acbp C. Candida albicans exhibits heterogeneous and adaptive cytoprotective responses to anti-fungal compounds. doi:10.1101/2022.07.20.500774

49. Diedrich J, Rehse SJ, Palchaudhuri S. Escherichia coli identification and strain discrimination using nanosecond laser-induced breakdown spectroscopy. Appl Phys Lett. 2007;90(16). doi:10.1063/1.2723659

50. Asadzadeh M, Ahmad S, Al-Sweih N, Khan Z. Rapid and Accurate Identification of Candida albicans and Candida dubliniensis by Real-Time PCR and Melting Curve Analysis. Med Princ Pract. 2018;27(6):543–548. doi:10.1159/000493426

51. Safavieh M, Coarsey C, Esiobu N, et al. Advances in *Candida* detection platforms for clinical and point-of-care applications. Crit Rev Biotechnol. 2017;37(4):441–458. doi:10.3109/07388551.2016.1167667

52. de Jong AW, Gerrits van den Ende B, Hagen F. Molecular Tools for Candida auris Identification and Typing. Methods Mol Biol. 2022;2517:33–41. doi:10.1007/978-1-0716-2417-3_3

53. Charlebois DA, Ribeiro AS, Lehmussola A, Lloyd-Price J, Yli-Harja O, Kauffman SA. Effects of Microarray Noise on Inference Efficiency of a Stochastic Model of Gene Networks.

54. Dingle TC, Butler-Wu SM. MALDI-TOF Mass Spectrometry for Microorganism Identification. Clin Lab Med. 2013;33(3):589–609. doi:10.1016/J.CLL.2013.03.001

55. Kurtzman CP, Robnett CJ. Identification of Clinically Important Ascomycetous Yeasts Based on Nucleotide Divergence in the 5 End of the Large-Subunit (26S) Ribosomal DNA Gene. Vol 35.; 1997. https://journals.asm.org/journal/jcm

56. Nucleotide BLAST: Search nucleotide databases using a nucleotide query. Accessed August 9, 2023. https://blast.ncbi.nlm.nih.gov/Blast.cgi?PROGRAM=blastn&BLAST_SPEC=GeoBlast&PAGE_TYPE=BlastSearch

57. Simonyan K, Zisserman A. Very Deep Convolutional Networks for Large-Scale Image Recognition. Published online September 4, 2014. http://arxiv.org/abs/1409.1556

58. Huang L, Qin J, Zhou Y, Zhu F, Liu L, Shao L. Normalization Techniques in Training DNNs: Methodology, Analysis and Application. IEEE Trans Pattern Anal Mach Intell. 2023;45(8):10173–10196. doi:10.1109/TPAMI.2023.3250241

59. Shorten C, Khoshgoftaar TM. A survey on Image Data Augmentation for Deep Learning. J Big Data. 2019;6(1). doi:10.1186/s40537-019-0197-0

60. classification.ipynb - Colaboratory. Accessed June 20, 2023. https://colab.research.google.com/github/tensorflow/docs/blob/master/site/en/tutorials/images/classification.ipynb#scrollTo=gN7G9GFmVr VY

61. Simonyan K, Zisserman A. Very Deep Convolutional Networks for Large-Scale Image Recognition.; 2014.

62. Szegedy C, Vanhoucke V, Ioffe S, Shlens J, Wojna Z. Rethinking the Inception Architecture for Computer Vision. Published online December 1, 2015.

63. He K, Zhang X, Ren S, Sun J. Deep residual learning for image recognition. In: *Proceedings of the IEEE Computer Society Conference on Computer Vision and Pattern Recognition*. Vol 2016-December. IEEE Computer Society; 2016:770–778. doi:10.1109/CVPR.2016.90

64. Tan M, Le Q V. EfficientNet: Rethinking Model Scaling for Convolutional Neural Networks. Published online May 28, 2019. http://arxiv.org/abs/1905.11946

65. Keras Applications. Accessed June 20, 2023. https://keras.io/api/applications/

66. Szegedy C, Vanhoucke V, Ioffe S, Shlens J, Wojna Z. Rethinking the Inception Architecture for Computer Vision. Published online December 1, 2015. http://arxiv.org/abs/1512.00567

67. Kumar R, Indrayan A. Receiver operating characteristic (ROC) curve for medical researchers. Indian Pediatr. 2011;48(4):277–287. doi:10.1007/s13312-011-0055-4

68. Yousif M, van Diest PJ, Laurinavicius A, et al. Artificial intelligence applied to breast pathology. Virchows Archiv. 2022;480(1):191–209. doi:10.1007/s00428-021-03213-3

69. Kurtzman CP, Fell JW, Boekhout T. The Yeasts : A Taxonomic Study. 5th ed. Elsevier; 2010. Accessed July 3, 2017. http://www.sciencedirect.com/science/book/9780444521491

70. Machuve D, Nwankwo E, Mduma N, Mbelwa J. Poultry diseases diagnostics models using deep learning. Front Artif Intell. 2022;5. doi:10.3389/frai.2022.733345

71. Ashkani-Esfahani S, Mojahed Yazdi R, Bhimani R, et al. Detection of ankle fractures using deep learning algorithms. Foot and Ankle Surgery. 2022;28(8):1259–1265. doi:10.1016/j.fas.2022.05.005

72. Cui X, Wei R, Gong L, et al. Assessing the effectiveness of artificial intelligence methods for melanoma: A retrospective review. J Am Acad Dermatol. 2019;81(5):1176–1180. doi:10.1016/j.jaad.2019.06.042

73. Geirhos R, Michaelis C, Wichmann FA, Rubisch P, Bethge M, Brendel W. Imagenet-trained CNNs are biased towards texture; increasing shape bias improves accuracy and robustness. 7th International Conference on Learning Representations, ICLR 2019. Published online 2019.

74. Heinke D, Wachman P, van Zoest W, Leek EC. A failure to learn object shape geometry: Implications for convolutional neural networks as plausible models of biological vision. Vision Res. 2021;189:81–92. doi:10.1016/J.VISRES.2021.09.004

75. Baker N, Lu H, Erlikhman G, Kellman PJ. Deep convolutional networks do not classify based on global object shape. PLoS Comput Biol. 2018;14(12). doi:10.1371/JOURNAL.PCBI.1006613

76. Pfaller MA, Diekema DJ. Role of Sentinel Surveillance of Candidemia: Trends in Species Distribution and Antifungal Susceptibility. J Clin Microbiol. 2002;40(10):3551–3557. doi:10.1128/JCM.40.10.3551-3557.2002

77. A. Shankarnarayan S, D. Guthrie J, A. Charlebois D. Machine Learning for Antimicrobial Resistance Research and Drug Development. In: The Global Antimicrobial Resistance Epidemic - Innovative Approaches and Cutting-Edge Solutions. IntechOpen; 2022. doi:10.5772/intechopen.104841

78. Wang Z, Guo D, Tu Z, et al. A Sparse Model-Inspired Deep Thresholding Network for Exponential Signal Reconstruction-- Application in Fast Biological Spectroscopy. IEEE Trans Neural Netw Learn Syst. Published online 2022:1–15. doi:10.1109/TNNLS.2022.3144580

79. Courtiol P, Maussion C, Moarii M, et al. Deep learning-based classification of mesothelioma improves prediction of patient outcome. Nat Med. 2019;25(10):1519–1525. doi:10.1038/s41591-019-0583-3

80. Shen S, Han SX, Aberle DR, Bui AA, Hsu W. An interpretable deep hierarchical semantic convolutional neural network for lung nodule malignancy classification. Expert Syst Appl. 2019;128:84–95. doi:10.1016/j.eswa.2019.01.048

81. Toda Y, Okura F. How Convolutional Neural Networks Diagnose Plant Disease. Plant Phenomics. 2019;2019. doi:10.34133/2019/9237136

82. Goebel R, Chander A, Holzinger K, et al. Explainable AI: The New 42? In: ; 2018:295–303. doi:10.1007/978-3-319-99740-7_21

83. Dhillon A, Verma GK. Convolutional neural network: a review of models, methodologies and applications to object detection. Progress in Artificial Intelligence. 2020;9(2):85–112. doi:10.1007/s13748-019-00203-0

84. Jensen J, Muñoz P, Guinea J, Rodríguez-Créixems M, Peláez T, Bouza E. Mixed fungemia: incidence, risk factors, and mortality in a general hospital. Clin Infect Dis. 2007;44(12):e109–14. doi:10.1086/518175

85. Li D, Wang Y, Hu W, et al. Application of Machine Learning Classifier to Candida auris Drug Resistance Analysis. Front Cell Infect Microbiol. 2021;11. doi:10.3389/fcimb.2021.742062

86. Bolin E, Lam W. A review of sensitivity, specificity, and likelihood ratios: evaluating the utility of the electrocardiogram as a screening tool in hypertrophic cardiomyopathy. Congenit Heart Dis. 2013;8(5):406–410. doi:10.1111/chd.12083

87. Classification: Precision and Recall | Machine Learning | Google for Developers. Accessed August 30, 2023. https://developers.google.com/machine-learning/crash-course/classification/precision-and-recall

88. Hicks SA, Strümke I, Thambawita V, et al. On evaluation metrics for medical applications of artificial intelligence. Sci Rep. 2022;12(1):5979. doi:10.1038/s41598-022-09954-8

89. Classification: ROC Curve and AUC | Machine Learning | Google for Developers. Accessed August 30, 2023. https://developers.google.com/machine-learning/crash-course/classification/roc-and-auc

